# Immortal genome assumption significantly underestimates replication and death rates of *Mycobacterium tuberculosis* in mice and monkeys

**DOI:** 10.1101/2025.08.15.670581

**Authors:** Allan D. Friesen, Vitaly V. Ganusov

## Abstract

Immune correlates of protection against infection with *Mycobacterium tuberculosis* (**Mtb**) or against tuberculosis (**TB**) remain poorly defined. The ratio of colony forming units (**CFUs**) to chromosomal equivalents (**CEQs**), *Z* = CFUs/CEQs, recovered either from the whole lung (mice) or individual lesions (monkeys or rabbits) of Mtb-infected animals has been used as a metric for how effectively Mtb is killed in vivo. However, the contribution of bacterial killing to changes in the CFU/CEQ ratio during an infection has not been rigorously investigated. We developed alternative mathematical models to study the dynamics of CFUs, CEQs, and their ratio during an Mtb infection. We find that the ratio *Z* alone cannot be used to infer the death rate of bacteria, unless the dynamics of CEQs and CFUs are entirely uncoupled, which is biologically unreasonable and inconsistent with the view that CEQs reflect an accumulated burden of both viable and non-viable bacteria. Importantly, we estimate a decay rate of 3.6%*/*day (a half life of about 20 days) of Mtb H37Rv CEQs in B6 mice that is similar to 4%*/*day, previously found for Mtb Erdman in cynomolgus macaques. While the estimated Mtb DNA decay rate seems small, we found that estimated rates of Mtb replication and death/killing are still extremely sensitive even to slow decay of Mtb DNA, in part, because Mtb replication and death rates are also small especially during chronic phases of infection. By applying our models to data on Mtb dynamics during the first 3 weeks of infection in macaques, we provide evidence of substantial killing of Mtb bacteria, prior to arrival of adaptive immunity to the site of infection, challenging the previously established notion of non-dying bacteria in the absence of T cell immunity and granuloma formation. We also propose experiments that will allow more accurately to measure the rate of Mtb DNA loss, helping more rigorously quantify impact of immunity on within-host Mtb dynamics.

## 1 Introduction

In many studies focused on identifying predictors of control of *Mycobacterium tuberculosis* (**Mtb**) during antibiotic treatment or following vaccination, colony forming units (**CFU**s) of Mtb isolated from whole lungs (mice) or from individual lung lesions (rabbits or macaques) are used as the primary metric for control of the bacteria population^1–5^. Efficacious treatment or vaccine-induced immune response may increase the bacterial clearance (death) rate, reduce the rate of Mtb replication, or impact both; but measurements of the total number of viable bacteria in a tissue typically do not allow to discriminate between these different effects^6–8^. Evaluating the impact of treatments and/or vaccination on Mtb replication or death rates requires development of additional metrics indicating how rapidly Mtb divides and dies in vivo^8,9^. The main focus of this paper is to determine how one new metric, the number of Mtb DNA molecules (chromosomal equivalents, **CEQs**), can be used to estimate the rates of Mtb replication and death in vivo.

In the first study to quantify replication and death rates of Mtb in vivo, Muñoz-Elías *et al*. ^6^ developed a rigorous methodology to measure CEQs as a metric for quantifying cumulative bacterial burden (**CBB**) during Mtb infection of mice. They found that CEQs in infected mice treated with isoniazid (**INH**) were stable over an 8 week treatment period, while CFUs declined by four orders of magnitude, concluding that Mtb CEQs are extremely stable in lungs of mice. They further used measurements of CFUs and CEQs over time in lungs of untreated Mtb-infected mice to conclude that Mtb replication is reduced by more 10 fold during the chronic phase (4-16 weeks after infection), compared with that during acute infection (first 2 weeks). This static picture of chronic Mtb infection in mice was challenged by Gill *et al*. ^10^ who used a “replication clock” plasmid to estimate Mtb replication and death rates in vivo^10,11^. This paper, along with subsequent analysis by McDaniel *et al*. ^12^, concluded that the Mtb replication rate declines only roughly 4-fold from acute to chronic infection.

Subsequent studies in non-human primates (**NHPs**) and rabbits used measurements of Mtb CFUs and CEQs to determine the replication state of Mtb in individual lung lesions (granulomas). In their pioneering study, Lin *et al*. ^13^ interpreted relatively stable CEQs per lesion between 4 and 11 weeks post-infection as indicating Mtb in a mostly non-replicating state in lungs of macaques. They furthermore introduced the CFU/CEQ ratio as a metric for “extent of killing” by the immune response. This metric has subsequently been used to evaluate the efficacy of different antibiotics in individual granulomas of rabbits^1,2,8,14,15^.

Although studies agree that with rise of adaptive (T cell) response in the lung, the rate of Mtb replication in the chronic infection is reduced, they disagree on the magnitude of this effect. Work with the replication clock plasmid suggests Mtb replication rate in chronic infection of mice and rabbits may be substantial^10,12,16^. In contrast, studies utilizing CEQs have generally reported stable CEQs during chronic infection in mice, monkeys, and rabbits, which has been interpreted as an indication that Mtb is non-replicating during chronic infection, under the assumption that CEQs are extremely stable (i.e., immortal). However, CEQ decline has been observed in chronically infected macaques, treated with INH, decaying at an rate of about 4%/day (see Supplemental Figure S2 in Lin *et al*. ^13^), and has also been reported in rabbits^1^. Whether such an apparently small genome decay rate is indeed negligible to infer the replicative state of bacteria in the chronic stage of infection has not been rigorously investigated.

Here we have developed mathematical models of within-host Mtb dynamics by explicitly tracking accumulation and loss of CFUs and CEQs. Our most general model describes several key processes in CFU and CEQ dynamics but it is over-parameterized for typical experimental data; we thus developed two alternative simplified versions of the general model that take different assumptions of how CEQs are produced during an infection. The independent dynamics (**ID**) model treats the dynamics of CFUs and CEQs independently, as in the Lin *et al*. ^13^ model, and the dependent dynamics (**DD**) model couples the two quantities by explicitly dividing the CEQs into contributions from culturable and dead bacteria, more similar to the approach of Muñoz-Elías *et al*. ^6^. Both models include the possibility of CEQ loss due to DNA degradation, absent from previous models. By reanalysis of the data on CEQ decay in INH-treated mice^6^ we estimate a loss rate of 3.6%/day, similar to the value found in macaques. We found that both ID and DD models can describe the CFU and CEQ data in mice or monkeys, suggesting that CFU and CEQ data alone are not sufficient to discriminate between the alternatives. Regardless of the model, accounting for CEQ loss results in substantially higher estimated replication and death rates of Mtb in chronic infection of mice (4-16 weeks post-infection) than that predicted by analyses assuming immortal CEQs. Furthermore, the DD model fitted to CFU and CEQ data in granulomas of macaques predicts substantial rates of both Mtb replication and death during the early phase of infection (first 3 weeks), prior to the arrival of T cell immunity in the lung. By using mathematical modeling and stochastic simulations we propose experiments that may help more precisely to estimate the CEQ decay rate in mice. Taken together, our mathematical modeling-based framework can be used to evaluate more rigorously the impact of vaccination and/or drug treatment on Mtb dynamics from CFU and CEQ measurements.

## 2 Materials & methods

### 2.1 Data

For our analyses we digitized data from two published studies^6,13^; these are typically averages per time point, since the data from individual animals were not published, and original data could not be located upon request. In the first experiment of Muñoz-Elías *et al*. ^6^, 24 mice were aerosol infected with approximately 200 CFU of Mtb (H37Rv or Erdman). Mice were sacrificed in groups of four at 1, 14, 28, 56, 84, and 112 days post-infection (or 0, 2, 4, 8, and 16 weeks post-infection). Lungs were harvested at necropsy, and CFUs and CEQs of Mtb in the lungs were measured (“dataset 1”, Figure 2A of Muñoz-Elías *et al*. ^6^). In the second experiment, 20 mice were inoculated intravenously with a dose ∼10^6^ CFU. Untreated mice were sacrificed at 4, 8, and 12 weeks post-infection. For the remaining mice, isoniazid (**INH**) was administered beginning at 4 weeks post-infection; four treated mice were sacrificed at 8 weeks and four more at 12 weeks post-infection. For all mice, CFUs and CEQs of Mtb in the lungs were measured (“dataset 2”, Figure 4A of Muñoz-Elías *et al*. ^6^).

In their study of Mtb Erdman dynamics in cynomolgus macaques, Lin *et al*. ^13^ infected macaques with a low dose (∼34 CFU) by bronchial instillation. Animals were sacrificed at 4 weeks (4 animals) and 11 weeks (3 animals). CFUs and CEQs of Mtb were measured from individual lesions from the lungs (“dataset 3”, Figure 3A&D of Lin *et al*. ^13^).

### 2.2 Mathematical models

#### Munoz-Elias et al. model

Muñoz-Elías *et al*. ^6^ proposed a discrete time-based mathematical model to describe the dynamics of CFUs and CEQs in their experiments. The model includes two parameters: a replication rate, *K*, and a net population growth rate, *K*^*′*^. The difference between *K*^*′*^ and *K* accounts for bacteria death. The model is simulated in discrete time steps; during each step, the viable population first grows exponentially with rate *K*, and then a portion of the bacteria are killed, so that the overall result is exponential growth with rate *K*^*′*^. The authors also assumed the CEQs do not decay. Although they mention that the model can be applied more generally, their analysis is restricted to the simple case of constant CFUs (*K*^*′*^ = 0) which results in linear growth of CEQs over time. Although their description of the model is somewhat unclear (e.g., no explicit equations were formally written), it seems conceptually similar to a discrete version of the dependent dynamics model we propose in this paper, for the special case of immortal genomes (see below).

#### Lin et al. model

Because in some experiments with Mtb in macaques, CEQs appear to be stable over time, Lin *et al*. ^13^ proposed a model in which CFUs (*B*) and CEQs (*Q*) are described independently by logistic models:

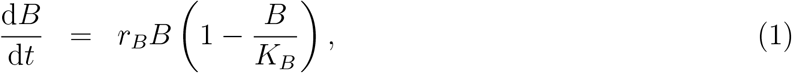

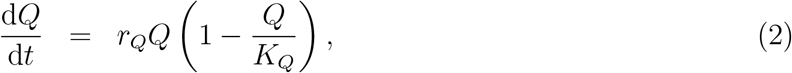

where *r*_*B*_ and *r*_*Q*_ are the net rates of replication (or death) of bacteria and genomes, respectively, and *K*_*B*_ and *K*_*Q*_ are the carrying capacity (saturation level) for bacteria and genomes, respectively. Different rates and carrying capacities are used at different stages of infection (*i*.*e*., in the first 4 weeks, and after 4 weeks). In the Supplement, we discuss details of this model and some difficulties in associating parameters of this model with actual physiological processes of Mtb replication and death in vivo. Despite not separating replication and death explicitly, the model has four parameters and two initial conditions, so that fitting the model to only CFU and CEQ data (2 measurements per time point) would typically result in overfitting and would require introducing additional assumptions to constrain model fits.

#### General model for CFU and CEQ dynamics

To track the dynamics of Mtb genomes we divide the Mtb population into culturable population *B* and a population that includes viable but not culturable on solid media (**VBNC**) bacteria and dead bacteria *D* (**Figure 1**A); both populations replicate (at rates *r*_*B*_ and *r*_*D*_, respectively) or die (at rates 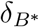 and *δ*_*D*_, respectively) and culturable bacteria may also convert into VBNC/dead state at rate *δ*_*B*_ (**Figure 1**A):

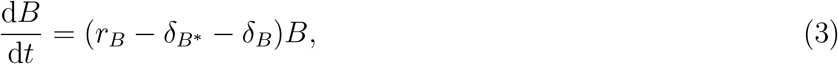

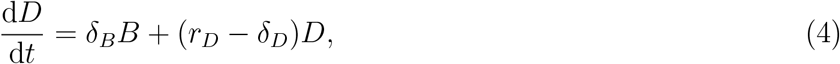

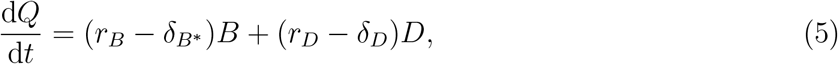

where the total number of Mtb chromosomal equivalents *Q* = *B* + *D*, and the death rate 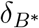 denotes death of culturable bacteria resulting in loss of the chromosome, e.g., due to degradation of the dead bacteria by macrophages. The general model has 7 parameters (5 rates and 2 initial conditions) that are not possible accurately to estimate from typical experimental data that has only 2 measurements (CFU and CEQ) per time point. Therefore, we consider the following alternative models that are extreme cases of a more general framework for modeling the dynamics of CFUs and CEQs.

#### Independent dynamics (ID) model

In the ID model, there are two independent quantities: the culturable population (CFUs), *B*, and the detectable chromosomal equivalents (CEQs), *Q* (**Figure 1**B). These populations reproduce with the same rate, *r*, but have different decay (death) rates, *δ*, and *δ*_*Q*_, respectively; so in the general model (**Figure 1**A) we set *r*_*B*_ = *r*_*D*_ = *r, δ*_*B*_ = 0, *δ*_*Q*_ = *δ*_*D*_, 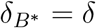, and *Q* = *D* resulting in

**Figure 1:**
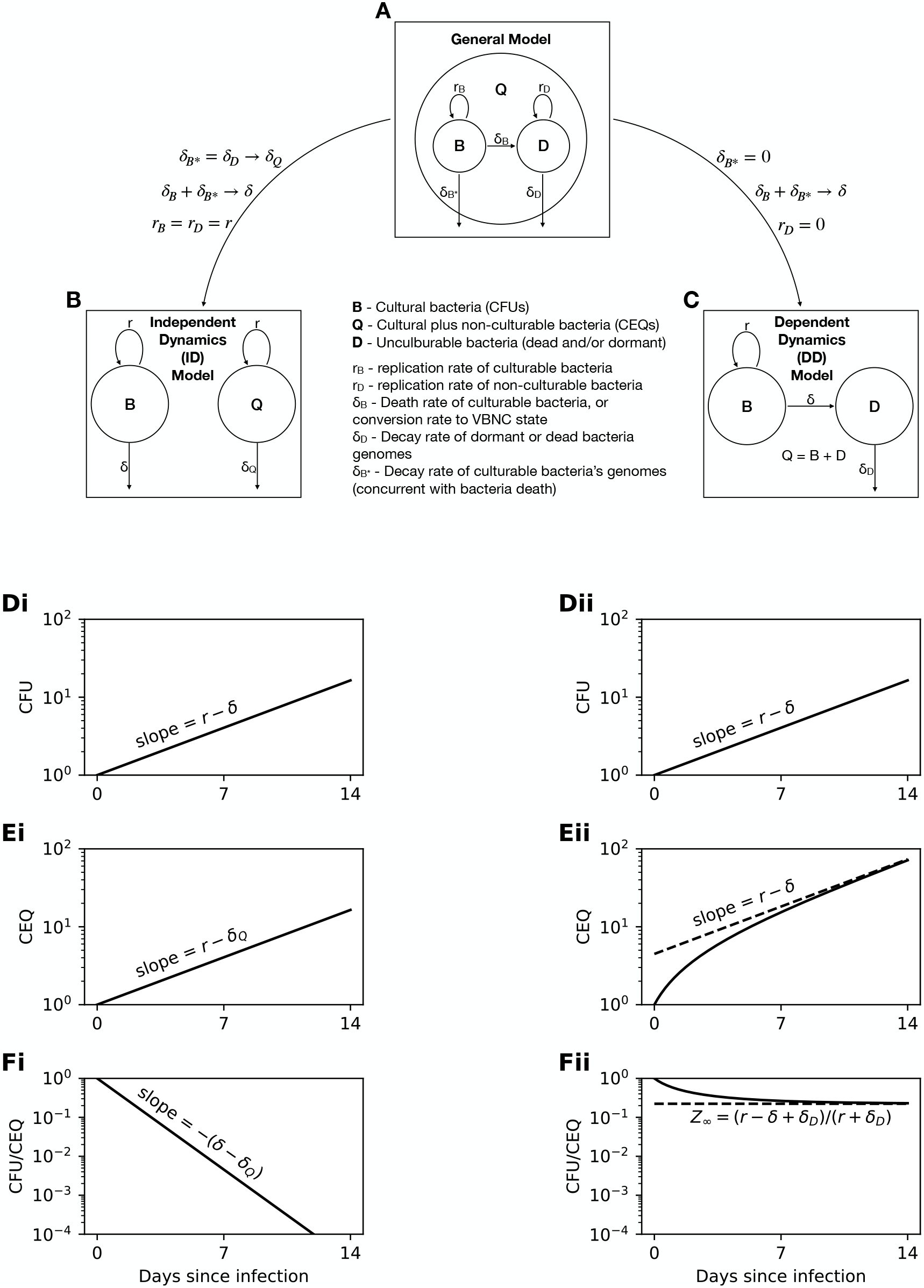
Framework for modeling dynamics of colony forming units (CFU) and chromosomal equivalents (CEQ) of Mtb in vivo. **A**: General model. Cultivable population *B* measurable as CFUs and nonculturable population *D* replicate and decay at rates (*r*_*B*_ and *r*_*D*_ or 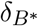 and *δ*_*D*_, respectively), and culturable bacteria convert to nonculturable bacteria with rate *δ*_*B*_. **B**: Independent dynamics (**ID**) model, in which the two populations replicate with rate *r*, but decay at different rates *δ* and *δ*_*Q*_. **C** Dependent dynamics (**DD**) model, in which all nonculturable bacteria are assumed to be dead and there is no loss of genomes due to killing of bacteria (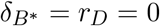). **D** – **F**: sketches of the dynamics of *B, Q*, and *Z*, predicted by the ID and DD models with constant replication and death rates.

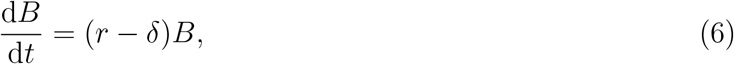

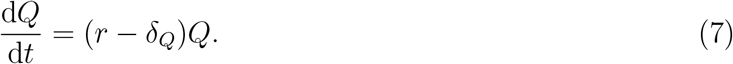

This model is equivalent to **eqns. (1)–(2)**, with appropriately chosen density-dependent *δ* and *δ*_*Q*_. For constant parameters, this model predicts exponential change in the total number of bacteria, genomes, or the ratio of bacteria to genomes (**Figure 1**Di-Ei-Fi).

#### Dependent dynamics (DD) model

In the DD model, we explicitly divide the total bacteria population (*Q* = *B* +*D*) into culturable (*B*) and dead (*D*) sub-populations (**Figure 1**C); we arrive at this model from the general model assuming 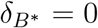, *δ*_*B*_ = *δ, r*_*B*_ = *r*, and *r*_*D*_ = 0. In the DD model, the viable population reproduces and dies with per capita rates *r* and *δ*, respectively, and the dead population does not reproduce and decays with per capita rate *δ*_*D*_ (resulting in the loss of Mtb genomes):

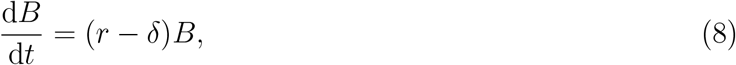

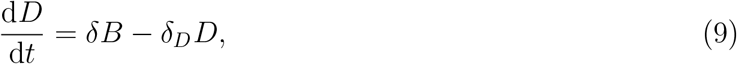

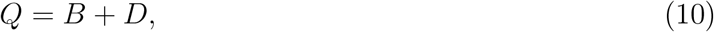

where *δ*_*D*_ is the CEQ decay rate. For constant parameters, this model predicts exponential change in the total number of bacteria but somewhat complex changes in the total number of genomes and of the ratio of bacteria to genomes (**Figure 1**Dii-Eii-Fii).

#### Flexible Independent dynamics (FID) model

To generate what we will call the flexible independent dynamics model (**FID**), we can consider the approximation of the general model (**eqns. (3)–(4)**) in which chromosomes of populations *B* and *D* replicate and die at different rates and 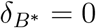:

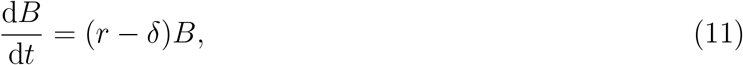

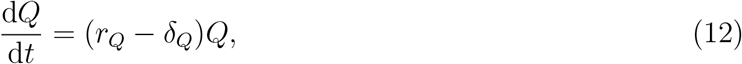

where we have renamed *r*_*B*_ = *r* and *δ*_*B*_ = *δ*. The FID model is equivalent to the ID model, and setting *r*_*Q*_ = *r* reduces the FID model to the ID model (**Figure 1**A-B). Similarly, by setting *δ*_*B*_ = *r*_*D*_ = 0 in **eqns. (3)–(4)** we will arrive to the DD model (**Figure 1**A&C).

#### Time-dependent replication and death rates

As in our previous analyses of dynamics of Mtb strains containing replication clock plasmid pBP10^12,16^, to accurately describe data on CFU and CEQ dynamics we assume that the rates of Mtb replication and death are constant in a given time period (defined by experimental measurements) but may change between time periods. Boundaries of these time periods depend on the study; for example, when fitting models to CFU/CEQ data in mice (Figure 2A in ref^6^, dataset 1) the replication and death rates are defined as follows:

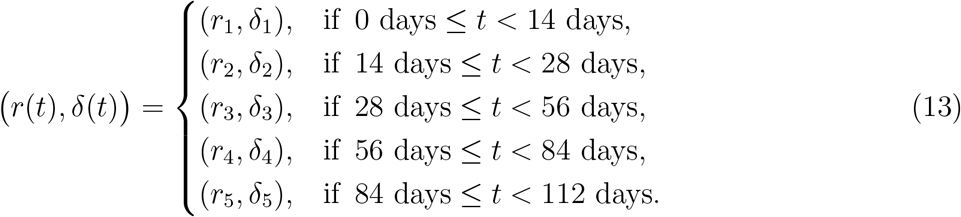

**Figure 2:**
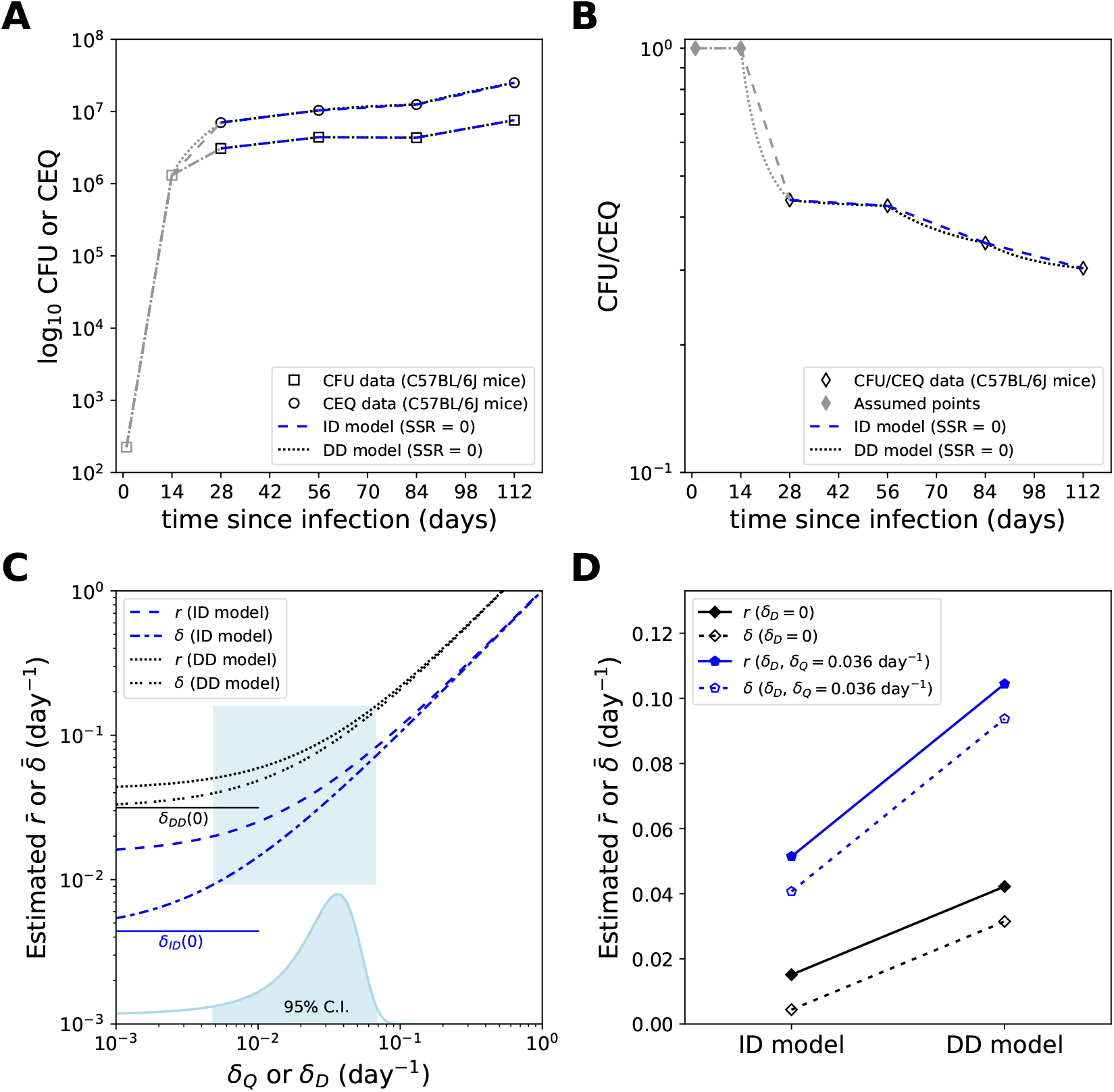
Estimates of Mtb replication (*r*) and death (*δ*) rates are strongly dependent on the decay rate of intact genomes independently of the underlying model of Mtb dynamics. **A-B**: Predictions of the ID or DD model fit to the CFU and CEQ data assuming genome decay rate of *δ*_*Q*_ = *δ*_*D*_ = 0.036*/*day. We show model fits to the CFU or CEQ data (**A**) or to the CFU/CEQ ratio (**B**). Parameters of the best fit are shown in **Table 1**. Note that we assumed particular values of the CFU/CEQ ratio *Z* during first 14 days of infection since these data were not available in the original publication ^6^. **C**: Dependence of estimated average replication 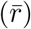 and death 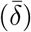 rates on the assumed value of genome decay rate (*δ*_*Q*_ or *δ*_*D*_ for ID or DD model, respectively). The distribution at the bottom of the graph is estimated from the decay rate of CEQs of Mtb in mice infected with Mtb, then treated with isoniazid (**Supplemental Figure S2**). The shown average rates are the mean of the three rates extracted from the three last time intervals shown in panels A and B (days 28 to 112, see **Table 1**). **D**: Comparison of average estimates of 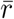 and 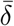 for the ID and DD models depending on the assumed genome decay rate (*δ*_*Q*_ or *δ*_*D*_ in ID or DD models, respectively).

**Table 1:** Parameters of the best fits of ID and DD models to CFUs and CEQs of Mtb in control/untreated mice. We fitted ID model (**eqns. (6)–(7)**) or DD model (**eqns. (8)–(10)**) to the data on CFU and CEQ dynamics in Mtb-infected mice ^6^ assuming that parameters are constant for a given time period (noted at the top row) but change between periods. Unfortunately, CEQs at 1 and 14 days post-infection were not measured in these experiments, therefore, we assumed that 1 day postinfection, when CFUs were measured, *Z*(1) = 1. We furthermore used different assumptions of the value of *Z* on day 14 to identify feasible ranges of *r* and *δ* during the first two time intervals. Minimum rates are obtained for the first time interval, when we assume *Z*(14) = 1. Maximum rates for the first time interval are obtained when *Z* takes a minimal value, which we take to be the larger of *Z*(28) or value of *Z*(14) that produces *r*_1_ = 1 day^−1^. We estimated the replication and death rates assuming two different values for the genome decay rates *δ*_*D*_ = 0.036*/*day or *δ*_*Q*_ = 0.036*/*day or both equal to 0.

| Model | $Z_{14}$ | $\delta_D$ or $\delta_Q$ ($\text{day}^{-1}$) | 1 - 14 days | | 14 - 28 days | | 28 - 56 days | | 56 - 84 days | | 84 - 112 days | |
| --- | --- | --- | --- | --- | --- | --- | --- | --- | --- | --- | --- | --- |
| | | | $r_1$ ($\text{day}^{-1}$) | $\delta_1$ ($\text{day}^{-1}$) | $r_2$ ($\text{day}^{-1}$) | $\delta_2$ ($\text{day}^{-1}$) | $r_3$ ($\text{day}^{-1}$) | $\delta_3$ ($\text{day}^{-1}$) | $r_4$ ($\text{day}^{-1}$) | $\delta_4$ ($\text{day}^{-1}$) | $r_5$ ($\text{day}^{-1}$) | $\delta_5$ ($\text{day}^{-1}$) |
| ID | 1 | 0.000 | 0.667 | 0.000 | 0.120 | 0.059 | 0.014 | 0.001 | 0.007 | 0.007 | 0.025 | 0.005 |
| ID | 0.439 | 0.000 | 0.731 | 0.063 | 0.061 | 0.000 | 0.014 | 0.001 | 0.007 | 0.007 | 0.025 | 0.005 |
| ID | 1 | 0.036 | 0.703 | 0.036 | 0.156 | 0.095 | 0.050 | 0.037 | 0.043 | 0.043 | 0.061 | 0.041 |
| ID | 0.439 | 0.036 | 0.767 | 0.099 | 0.097 | 0.036 | 0.050 | 0.037 | 0.043 | 0.043 | 0.061 | 0.041 |
| DD | 1 | 0.000 | 0.667 | 0.000 | 0.197 | 0.136 | 0.032 | 0.019 | 0.017 | 0.018 | 0.077 | 0.057 |
| DD | 0.679 | 0.000 | 1.000 | 0.333 | 0.174 | 0.113 | 0.032 | 0.019 | 0.017 | 0.018 | 0.077 | 0.057 |
| DD | 1 | 0.036 | 0.667 | 0.000 | 0.228 | 0.167 | 0.080 | 0.067 | 0.077 | 0.078 | 0.155 | 0.135 |
| DD | 0.667 | 0.036 | 1.000 | 0.333 | 0.212 | 0.151 | 0.080 | 0.067 | 0.077 | 0.078 | 0.155 | 0.135 |

It is sometimes useful to coarse grain the infection into an “early” or “acute” stage, during which Mtb number in the lung grows approximately exponentially, and a “chronic” stage, during which Mtb number is approximately stable. For purposes of this paper, acute or early infection refers to the first 2 weeks in mice and the first 3 - 4 weeks in macaques. Mtb-infected mice settle to a chronic infection by approximately 8 weeks post-infection. In this paper, when we calculate mean replication and death rates for chronic infection in mice (**Figure 2**C-D), we also include the rate between 4 and 8 weeks post-infection, since in the Muñoz-Elías *et al*. ^6^ experiment, the changes in CFU and CEQ levels during this period appear consistent with later time intervals. We also report estimates of Mtb replication and death rates for individual time intervals (**Table 1**). Note that in these analyses we assume that the CEQ decay rate (i.e., *δ*_*Q*_ in ID model or *δ*_*D*_ in DD model) does not depend on time since infection.

### 2.3 Power Analysis

We performed a power analysis for detection of the decay rate *δ*_*D*_ of detectable Mtb genomes in mice treated with antibiotics. We assumed the dynamics follow the DD model. We used digitized data from Muñoz-Elías *et al*. ^6^ (“dataset 2”) to estimate the CFUs of Mtb in the lungs of mice 4 weeks after infection. Since the DD model does not make mechanistic sense when CEQs are less than the CFUs, we used average values of CEQs such that the average of *Z* was 0.9, 0.5, or 0.1. For each case, we drew values of *D* = *Q* − *B* from a log-normal distribution and added this to *B* to obtain initial values of *Q*. We used the published calibration curve^6^ to estimate the uncertainty in *Q* as approximately 0.6 logs; this may be a somewhat pessimistic estimate, since measurements of *Q* sometimes appear to follow tighter distributions. However, we prefer to err toward caution when estimating statistical power. We chose the width of the log-normal distribution of *D* such that it reproduces this uncertainty. After drawing initial values from these distributions, we integrated the DD model and evaluated the best fit line of ln *Q* vs. *t* between two chosen time points. This process was repeated for each mouse that would be needed in the proposed experiment; the whole process was then repeated 10,000 times for each set of parameters (each number of mice, each *δ*_*D*_, *etc*). An F-test for nested models was used (SciPy function stats.f.cdf) to obtain a *p* value for each simulated experiment, where the full model assumes that genomes decay (*δ*_*D*_ *>* 0) and the reduced model assumes that genomes do not decay (*δ*_*D*_ = 0). We used *p* = 0.05 as a cut-off for statistical significance that detected *δ*_*D*_ *>* 0.

### 2.4 Quantifying uncertainty in estimated genome decay rate

To estimate the distribution of possible decay rates of intact genomes, we digitized mean values and error bars of *Q* from the final two points of Fig 4A in Muñoz-Elías *et al*. ^6^. The reported error bars are based on standard deviations of the absolute number (not logarithms) of CEQs. For points at 8 and 12 weeks post-infection, we used the digitized mean and standard deviation to define the corresponding log-normal distribution that has the same mean and standard deviation, using standard formulas^17^. We then propagated this quantity to the error in the genome decay rate, 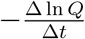 measured between 8 and 12 weeks post-infection in INH-treated mice assuming normal distribution of the error (**Figure 2**C). The uncertainty results from the numerator, which is a difference of the two quantities. Because measurements at 8 and 12 weeks post-infection are expected to be statistically independent, the distribution of the difference Δ ln *Q* follows a lognormal distribution with variance given by the sum of the squared standard errors at 8 and 12 weeks post-infection. We also estimated *δ*_*D*_ by fitting the DD model with a single set of parameters to the mean CFUs and CEQs from Fig 4A in Muñoz-Elías *et al*. ^6^ (**Supplemental Figure S2**). The resulting estimate of *δ*_*D*_ was very close (0.035 day^−1^) to the mean of the estimated distribution (0.036 day^−1^) estimated from the slope of ln *Q* between 8 and 12 weeks post-infection in INH-treated mice. Note that the genome decay rate cannot be estimated if the dynamics of *B* and *Q* are treated independently (e.g., in the ID model), since the dynamics of *Q* will reflect a combination of *r* and *δ*_*Q*_, even when *B* is much less than *Q* (see **eqn. (7)**).

### 2.5 Stochastic Simulations

We performed stochastic simulations of Mtb dynamics with an adaptive tau leaping algorithm^18^ that reduces to the full stochastic simulation algorithm (**SSA**)^19^ when populations are small. We implemented this algorithm in Fortran 08 for increased flexibility and computation speed with a wrapper in python, but compared some test cases to the Gillespy2 python package, as a point of caution.

## 3 Results

### Models of dynamics of CFUs and CEQs

When following Mtb dynamics in infected animals, it is typical to measure the number of viable bacteria (bacteria that are able to grow on plates, i.e., CFUs) in a given tissue. More recently, a new metric, the total number of Mtb DNA molecules or chromosomal equivalents (**CEQs**), has been used as measure of cumulative bacterial burden (**CBB**). CEQ measurements have been used to gain additional insights into details of in-host Mtb dynamics, e.g., impact of various antibiotics on rates of Mtb replication and death in granulomas of rabbits^1,2,6,13^. Despite its use, however, interpretation of data on Mtb CEQ dynamics has been semi-quantitative, typically assuming that CEQs decay very slowly (or are immortal). We therefore sought to derive mathematical models that would describe generation of CEQs during an in vivo infection and apply these models to estimate the CEQ decay rate and how quickly Mtb replicates and dies in vivo.

CEQs are the total number of Mtb chromosomes found in a tissue and are expected to be calculated from live (platable), dead, and viable-but-not-culturable (**VBNC**) bacteria. Therefore, the general model of Mtb dynamics that accounts for CEQs should include these sub-populations. In our general model, however, we started with just two of such populations – viable bacteria *B* and dead/VBNC bacteria *D* (**Figure 1**A and **eqns. (3)–(4)**). Both populations can replicate and die at different rates, and viable bacteria can also convert into dead/VBNC bacteria (at rate *δ*_*B*_). Then the CEQs are given by the total number of chromosomes *Q* = *B* + *D* (**Figure 1**A), and loss of CEQs in this model is determined by the rates 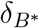 and *δ*_*D*_. While being fairly simplistic (e.g., the model lumps together dead and VBNC bacteria) this model in total has 7 parameters (5 rates and 2 initial conditions) that would not be possible to estimate from just two experimental measurements of CFUs and CEQs (per time point) given that model parameters are also likely to change with time since infection^12^. Therefore, we focused on two alternative simplifications of the general model: independent dynamics (**ID**) and dependent dynamics (**DD**) models (**Figure 1**).

In the ID model we assume that CFUs and CEQs are independent of each other (i.e., *δ*_*B*_ = 0 in the general model, **Figure 1**A&B); CFUs and CEQs replicate with the same per capita rate *r*, but decay with different rates *δ* (CFUs) and *δ*_*Q*_ (CEQs) (**eqns. (6)–(7), Figure 1**B). The ID model (and its extension, flexible independent dynamics (**FID**) model, **eqns. (11)–(12)**) are similar to the model proposed by Lin *et al*. ^13^ to describe the dynamics of CFUs and CEQs in granulomas of Mtb-infected monkeys. In the ID model, when rates are constant the dynamics of CFUs (*B*), CEQs (*Q*), and the ratio CFUs/CEQs (*Z*) are each described by exponential growth or decay (**Figure 1**Di–Fi). Importantly, in this model, the decay rate of the CFU/CEQ ratio does not depend on the replication rate *r* and is determined only by the difference *δ* − *δ*_*Q*_ (**Figure 1**B and **eqn. (S.7)**). Therefore, if the CEQ decay rate is known, the death rate *δ* in the ID model can be uniquely determined from the dynamics of the CFU/CEQ ratio. However, there are conceptual difficulties with associating *δ* of this model with the actual death rate of viable bacteria (see Supplemental Information for more detail).

In contrast, in the DD model we assume that upon death viable bacteria *B* become dead bacteria *D* (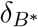 and *δ* = *δ*_*B*_ in the general model) that do not replicate (*r*_*D*_ = 0 in the general model) and DNA in the dead bacteria decays over time (at rate *δ*_*D*_, see **eqns. (8)– (10)** and **Figure 1**C). In the DD model, the total CEQs, *Q*, is then simply the sum of CFUs *B* and detectable genomes of dead bacteria *D*; that is, *Q* = *B* + *D*. This model is similar to that described verbally in Muñoz-Elías *et al*. ^6^. In the DD model when rates are constant CFUs grow or decline exponentially over time (**Figure 1**Dii); however, the dynamics of the CEQs and the CFU/CEQ ratio are more complex (**Figure 1**Eii&Fii). In particular, the CFU/CEQ ratio asymptotically approaches a limiting value (carrying capacity) when bacterial burdens are increasing (**Figure 1**Fii and **eqn. (S.10)**), but declines exponentially when bacterial burdens are decreasing (**Supplemental Figure S1** and **eqn. (S.14)**). Taken together, the dynamics of CEQs and CFU/CEQ ratio are different between ID and DD models allowing, at least in principle, to discriminate between these alternatives using experimental data.

### Estimating CEQ decay rate

Our analysis of the alternative ID and DD models indicates that dynamics of the CFU/CEQ ratio depends on the CEQ decay rate (**Figure 1**F). Furthermore, in both models a change in CFUs, CEQs, or CFU/CEQ ratio between two time points depends on three parameters (e.g., *r, δ*, and *δ*_*D*_ in the DD model, see **Figure 1**Dii-Fii), so that knowing the rate of CEQ decay is necessary to estimate the rates of Mtb replication and death. We therefore digitized and analyzed data from a previous experiment^6^ in which mice were infected intravenously with 10^6^ CFUs of Mtb and then treated with INH starting 28 days post-infection for 56 days (**Supplemental Figure S2**A). As expected, the number of viable bacteria declined exponentially (at a rate 0.17*/*day); interestingly, the CEQ number increased in the first 28 days during treatment and then declined (**Supplemental Figure S2**A). Because the number of viable bacteria was relatively small at 56 days post-infection (28 days post-treatment), CEQs can be considered independent of CFUs between 56 and 84 days post-infection. By using linear regression we found that CEQs decline (between days 56 and 84) at a rate *δ*_*Q*_ = *δ*_*D*_ = 0.036*/*day that is the estimated CEQ decay rate.

The increase and then decline in CEQs observed in INH-treated mice between 28-56 and 56-84 days post-infection (**Supplemental Figure S2**A) are not fully consistent with predictions of the ID model unless the CEQ replication rate is high initially and declines after day 56. In contrast, the DD model theoretically is able to predict increase in CEQ numbers as viable bacteria replicate and die. Indeed, we found that the DD model can well describe both CFU and CEQ data in INH-treated mice (**Supplemental Figure S2**A); the fits predicted CEQ decay rate of *δ*_*D*_ = 0.035*/*day consistent with the simple regression analysis (see above). However, to explain a relatively large initial increase in CEQs during the treatment (days 28-56 post infection) we found Mtb must be replicating and dying at relatively high rates (*r* = 1.29*/*day and *δ* = 1.45*/*day, **Supplemental Figure S2**A). Taken together, our analysis of the CEQ dynamics in INH-treated mice^6^ suggests that Mtb genomes have a non-zero decline rate of *δ*_*D*_ = 0.036*/*day; this is similar to the value estimated for Mtb Erdman in macaques ^13^.

### ID and DD models accurately describe the dynamics of CFUs and CEQs of Mtb in mice

Having estimated the CEQ decay rate in mice, we next sought to determine how well our alternative (ID and DD) models may fit the data on Mtb dynamics in mice. We therefore digitized data from experiments of Muñoz-Elías *et al*. ^6^ who aerosol-infected 24 mice with a standard dose of Mtb and measured Mtb CFUs and CEQs in the lungs over time (**Figure 2**A&B). We should note that CEQ measurements were not available at day 1 and 14 in the original paper^6^; therefore, when fitting models to data we initially made the assumption that *Z* = 1 at both times (**Figure 2**B).

Interestingly, we found that both ID and DD models can describe the data with reasonable parameter values and the assumed decay rate of Mtb genomes of 0.036 day^−1^; in fact, both models can fit the (averaged) data perfectly, with the sum of squared residuals (**SSR**) equal to 0. Because CEQs were not available for days 1 and 14, in the fits we assumed that the CFU/CEQ ratio *Z* = 1 at one day post-infection. In order to bracket possible rates, we then alternately assumed either that the CFU/CEQ ratio was also 1 on day 14, or that the CFU/CEQ ratio on day 14 equaled that on day 28. Because we fit with new parameters for each time interval, and because we fit to averages (individual mouse data were unavailable), both models fit the data again with SSR = 0.

Fitting two models with two assumed genome decay rates (i.e., *δ*_*D*_ = *δ*_*Q*_ = 0 or 0.036/day) and two assumed values of CFU/CEQ ratio *Z* on day 14, resulted in a total of 8 fits to this dataset. We found that in all cases, we could fit the data with parameters in a reasonable range (replication rates ≤ 1*/*day and all rates *>* 0, **Table 1**). Interestingly, we also could accurately fit the DD model to data from an independent experiment of intravenous infection of mice with Mtb (**Supplemental Figure S2**B).

The observation that either ID or DD model can fit the data well points to limitation of CFU/CEQ data alone to determine the underlying mechanisms behind the observed dynamics. Considered biologically, the ID and DD models make different assumptions of how CEQs are produced during the infection. The DD model assumes a mechanism in line with the language typically used^6,13^, referring to viable/culturable and dead bacteria. However, replication of Mtb is known to be heterogeneous, so that the DD model, in the form considered here, is somewhat simplistic. On the contrary, the ID model, despite mathematical simplicity, is challenging to interpret biologically. As a limit of the general model, the ID model assumes that the apparent death of platable bacteria is actually a transition to a VBNC state, in which bacteria continue to replicate with the same rate (see Supplemental Information for a more detailed discussion). Given the ability to describe the same dynamics with such contrasting pictures, more information is required in order to discriminate between possible interpretations of the data.

### Estimated replication and death rates depend strongly on the assumed decay rate of detectable Mtb genomes

Our analysis of the ID and DD models suggests that estimated replication and death rates are strongly sensitive to the decay rate of detectable genomes (e.g., see **eqns. (S.16)–(S.18)**); interestingly, however, we found that both ID and DD models could fit the CFU/CEQ data with excellent quality (SSR = 0) assuming immortal genomes (**Figure 2**A&B) but with different estimates of the Mtb replication and death rates (**Table 1**).

To more systematically investigate the impact of the assumed genome decay rate on Mtb replication and death rates, we varied the genome decay rate in the range 0.001 − 0.1 day^−1^ (the 95% confidence interval of the genome decay rate estimated in this paper) and estimated average replication (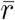) and death (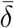) rates in the ID or DD models (**Figure 2**C), during chronic infection in mice (4 to 16 weeks post-infection). According to both models, higher genome decay rates result in higher estimated average replication and death rates (**Figure 2**C), and parameters of the ID model more strongly depend on the genome decay rate. Estimated rates of Mtb replication and death are about 2-3 fold larger for a decay rate of 0.036/day, compared with estimates obtained assuming immortal Mtb genomes. Given the imprecision in the estimated genome decay rate, actual rates of Mtb replication and death may be yet another 2-3 fold larger as compared to values obtained assuming *δ*_*D*_ = 0.036*/*day. At larger values of the genome decay rate estimated replication and death rates scale approximately linearly with *δ*_*D*_ or *δ*_*Q*_ (**Figure 2**C).

In addition to dependence on the genome decay rate, estimated replication and death rates depend on the model used to fit the data. In particular, for the same assumed genome decay rate we find about twice higher rates of Mtb replication and death in the DD model as compared to those in the ID model (**Figure 2**D). It should be noted, however, that direct comparison of estimated replication and death rates in the two models is not fully appropriate since the parameters *r* and *δ* have different interpretations in the different models. Nevertheless, considered at face value, independent dynamics of CFUs and CEQs leads to substantially smaller estimates of replication and death rates. When the decay rate of detectable genomes is assumed to be 0, there is an even larger discrepancy in 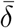, between the two models (**Figure 2**D).

### Both replication and death contribute significantly to Mtb dynamics during acute infection in macaques

Our analysis of the two alternative models of CFU and CEQ dynamics suggests that estimates of the Mtb replication and death rates strongly depend on the assumed longevity of Mtb genomes (e.g., see **eqns. (S.16)–(S.19)**). Importantly, given the estimated CEQ decay rate of *δ*_*D*_ = 0.036*/*day we found that Mtb replicates and dies at relatively high rates during chronic infection (4-16 weeks) of mice thus challenging the previously held view of “static equilibrium” (**Figure 2, Table 1** and ref.^6^). We therefore next sought to investigate whether the changing the assumption of immortal Mtb genomes may change interpretation of data on Mtb dynamics in acute (first 3 weeks) infection in macaques. Because we found that CFU and CEQ data in mice were insufficient to discriminate between ID and DD models, here, we start the data analysis with the DD model because this model more naturally allows to explain the rise in CEQ numbers during the first weeks of infection.

Lin *et al*. ^13^ were first to comprehensively follow dynamics of CFUs and CEQs in individual granulomas of macaques infected with Mtb Erdman. For the first 3-4 weeks after infection there was a rapid increase in the average number of CFUs and CEQs in the granulomas; interestingly, while the average CFU per granuloma declined after week 4, CEQ per granuloma remained relatively constant. These data were interpreted that in the first 3 weeks, prior to arrival of T cell response to the lung, Mtb replicates with minimal death but after 4 weeks, replication is halted (no change in CEQ numbers) and bacteria are being eliminated by the immune response^13^. We therefore investigated whether such interpretation is correct given that in mice and in monkeys CEQs have appreciable decay rate (*δ*_*D*_ = 0.036 − 0.04*/*day).

Because we did not have access to original CFU and CEQ data, we digitized median CFU and CEQ loads of individual lesions from Figure 3A&D of Lin *et al*. ^13^. Similar to the dataset on Mtb infection in mice^6^, CEQs were not available at 3 weeks post-infection. Therefore, we again fit the DD model twice, with different assumed initial rates assuming *δ*_*D*_ = 0 day^−1^ or *δ*_*D*_ = 0.036 day^−1^ and variable early replication and death rates.

We found that independently of the genome decay rate, the DD model with different Mtb replication and death rates could accurately fit the data; in particular, a model assuming that there is no death of bacteria (*δ* = 0) or a model with significant early death (*δ* ≈ 0.6*/*day), fitted the data with similar quality (**Figure 3**A&B, SSR = 0 in both cases). However, these two extreme fits predicted different changes in the rate of Mtb replication in the first 4 weeks of infection. For fits of the DD model to the data assuming that *δ* = 0 during the first 3 weeks of infection, a dramatic increase in Mtb replication rate *r* must occur around and prior to the onset of adaptive immunity (3 weeks), in order to fit the observed changes in CFU and CFU/CEQ ratio (**Figure 3**B&C and **Table 2**). This model prediction occurs because at 3 weeks post-infection, there has to be a large increase in Mtb replication rate (to generate CEQs) and a corresponding increase in the death rate (to avoid an increasing net growth rate of CFUs).

**Figure 3:**
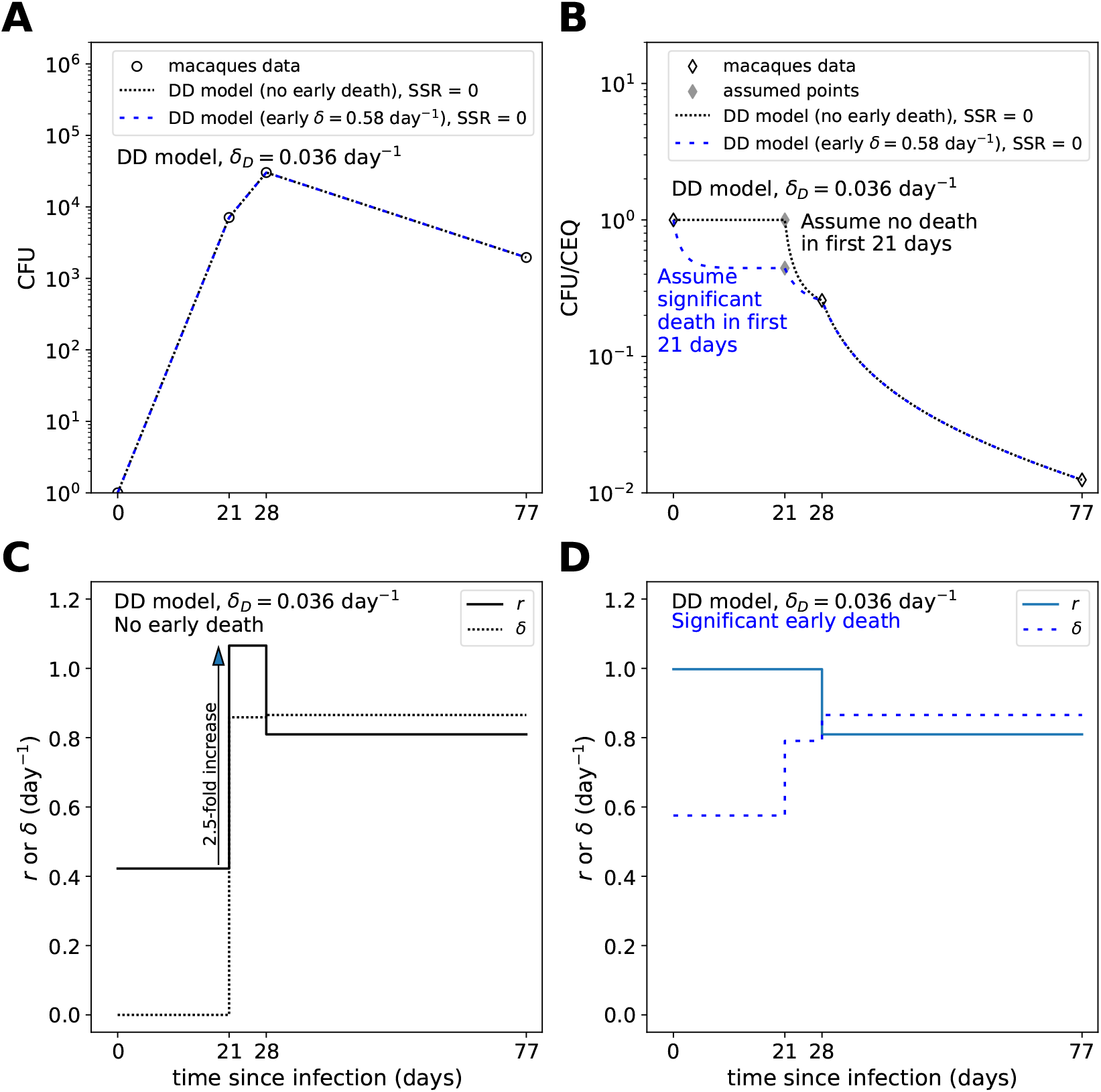
Both replication and death rates contribute significantly to Mtb dynamics in monkeys during acute infection. We digitized the data on CFU and CEQ dynamics from Lin *et al*. ^13^ and fitted alternative models to these data. **A**: Fits of the DD model to CFUs and CEQs data from Mtb-infected macaques ^13^. Symbols *r*_0_ and *δ*_0_ refer to model parameters during the acute phase of infection (day 0 to 21). **B**: Predictions of the DD model for CFU/CEQ ratio. Because CEQs at 21 days post infection were not given in these experiments, we assumed two extreme cases of parameters during the first 21 days: no death in the first 21 days, and maximum replication (taken as *r*_0_ = 1 day^−1^) and estimated associated death rate. These two scenarios generated different predictions on CFU/CEQ ratio dynamics. **C**: DD model-based predictions of the changes in the rates of Mtb replication (*r*) and death (*δ*) over time, assuming no Mtb death during first 21 days post-infection. The arrow shows a 2.5-fold increase in the predicted rate of Mtb replication between 21 and 28 days post-infection. **D**: DD model-based predictions of the changes in the rates of Mtb replication (*r*) and death (*δ*) over time, assuming substantial Mtb death during first 21 days post-infection. Under this assumption, *r* decreases and *δ* increases when adaptive immunity sets in, though the two rates remain similar in size and large in comparison with the net rate of decline *δ* − *r*. In fits we assumed *δ*_*D*_ = 0.036 day^−1^ as estimated for mice (**Supplemental Figure S2**) or monkeys ^13^. Other parameters for the fits as well as fits of the ID model and fits with *δ*_*D*_ = 0 or *δ*_*Q*_ = 0 are shows in **Table 2**.

**Table 2:**
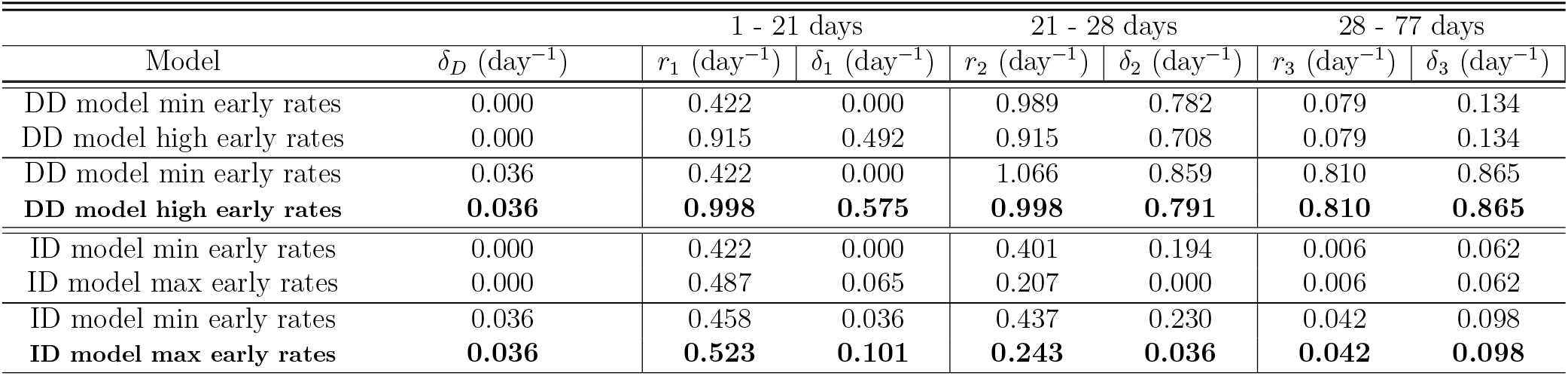
Parameters of fits of ID and DD models to Mtb (Erdman) CFU and CEQ data in lesions of macaques. We digitized the data on CFU and CEQ dynamics from Lin *et al*. ^13^ and fitted alternative models to these data (see **Figure 3**). Because CEQs at 21 days post-infection were not shown in original publication, we considered extreme assumptions to generate a range of feasible possibilities of estimated parameters during early infection. Minimum early rates are obtained by assuming *Z* = 1 on day 21 of infection, and maximum early rates are obtained by setting *Z* at 21 days post-infection equal to *Z* at 28 days post-infection. In some cases the latter assumption led to replication rates significantly in excess of 1 day^−1^. In these cases, we set *r* ≈ 1*/*day. It turned out that setting *r* = 1*/*day in some cases led to very small differences in rate between weeks 1-3 and week 4 of infection. In these cases, we fit the data with *r* kept constant during the first 4 weeks of infection.

| Model | $\delta_D$ (day<sup>-1</sup>) | 1 - 21 days | | 21 - 28 days | | 28 - 77 days | |
| --- | --- | --- | --- | --- | --- | --- | --- |
| | | $r_1$ (day<sup>-1</sup>) | $\delta_1$ (day<sup>-1</sup>) | $r_2$ (day<sup>-1</sup>) | $\delta_2$ (day<sup>-1</sup>) | $r_3$ (day<sup>-1</sup>) | $\delta_3$ (day<sup>-1</sup>) |
| DD model min early rates | 0.000 | 0.422 | 0.000 | 0.989 | 0.782 | 0.079 | 0.134 |
| DD model high early rates | 0.000 | 0.915 | 0.492 | 0.915 | 0.708 | 0.079 | 0.134 |
| DD model min early rates | 0.036 | 0.422 | 0.000 | 1.066 | 0.859 | 0.810 | 0.865 |
| **DD model high early rates** | **0.036** | **0.998** | **0.575** | **0.998** | **0.791** | **0.810** | **0.865** |
| ID model min early rates | 0.000 | 0.422 | 0.000 | 0.401 | 0.194 | 0.006 | 0.062 |
| ID model max early rates | 0.000 | 0.487 | 0.065 | 0.207 | 0.000 | 0.006 | 0.062 |
| ID model min early rates | 0.036 | 0.458 | 0.036 | 0.437 | 0.230 | 0.042 | 0.098 |
| **ID model max early rates** | **0.036** | **0.523** | **0.101** | **0.243** | **0.036** | **0.042** | **0.098** |

Alternatively, fitting the DD model with the opposite extreme assumption, that *r* and *δ* are both large during the early infection, leads to a picture in which both *δ* and *r* contribute significantly to the internal population dynamics, both during the first 3 weeks of infection and after the onset of adaptive immunity, and while the Mtb death rate increases over time, Mtb replication rate moderately decreases over time (**Figure 3**B&D, and **Table 2**). Because it is very difficult to envision rapid increase in Mtb replication rate between 3 and 4 weeks post-infection (**Figure 3**C) a model in which there is little Mtb death prior to arrival of adaptive immunity to the lung is unlikely. Therefore, a model in which there is rapid Mtb replication and substantial Mtb death prior to T cell immunity, in the first 3 weeks after infection (**Figure 3**D) is more consistent with the data.

While the DD model seems more reasonable to explain early accumulation of Mtb genomes, we nevertheless investigated whether the ID model may deliver different conclusions about early Mtb replication and death rates. As in the case of Mtb dynamics in mice, the ID model could fit the data on Mtb dynamics in macaques with excellent quality and reasonable parameter estimates (i,e., replication rates ≤ 1.07 per day, and all rates ≥ 0) (**Table 2**). However, the ID model predicted much lower Mtb replication rates, with the largest effect occurring during after 4 weeks infection (the effect on replication rate ranges from a factor of about 2 to a factor of about 20, **Table 2**). Thus, while both ID and DD models could accurately describe the CFU and CEQ data in individual granulomas of NHPs, they provided different estimates of Mtb replication and death rates.

### Power analysis to determine decay rate of Mtb genomes

Given the dependence of the Mtb genome decay rate on inferred Mtb replication and death rates both in mice and monkeys (**Tables 1 and 2**), we sought to investigate different types of experimental designs that would allow more accurately estimate the genome decay rate. We focus our analysis on Mtb dynamics in mice but similar arguments may be applied to studies with Mtb in rabbits or NHPs.

To accurately quantify the decay kinetics of Mtb genomes one must uncouple the dynamics of CFU and CEQs by using, for example, antibiotic treatment (e.g., **Supplemental Figure S2**). In one such experimental design we allow for Mtb replication in mice for 28 days (to allow for accumulation of a sufficient number of Mtb genomes) and then start efficient antibiotic treatment (**Figure 4**A). Previous experiments suggest that because of continuous accumulation of Mtb genomes during initial phase of treatment (either as dead or VBNC bacteria), it is important to measure CFUs and CEQs at some intermediate time point, e.g., at 56 days post-infection. Then, after additional 28 days we do a final measure of CFU and CEQ numbers (**Figure 4**A). In total, this would require 3*n* mice per experiment with *n* mice sampled at each time point. The genome decay rate is then evaluated as the slope of − ln *Q* vs. time between the 56 and 84 days post-infection (**Figure 4**A). While measuring CFUs and CEQs at 28 days post-infection is not strictly necessary to estimate the genome decay rate, knowing these numbers will help paint a more complete picture of Mtb dynamics.

**Figure 4:**
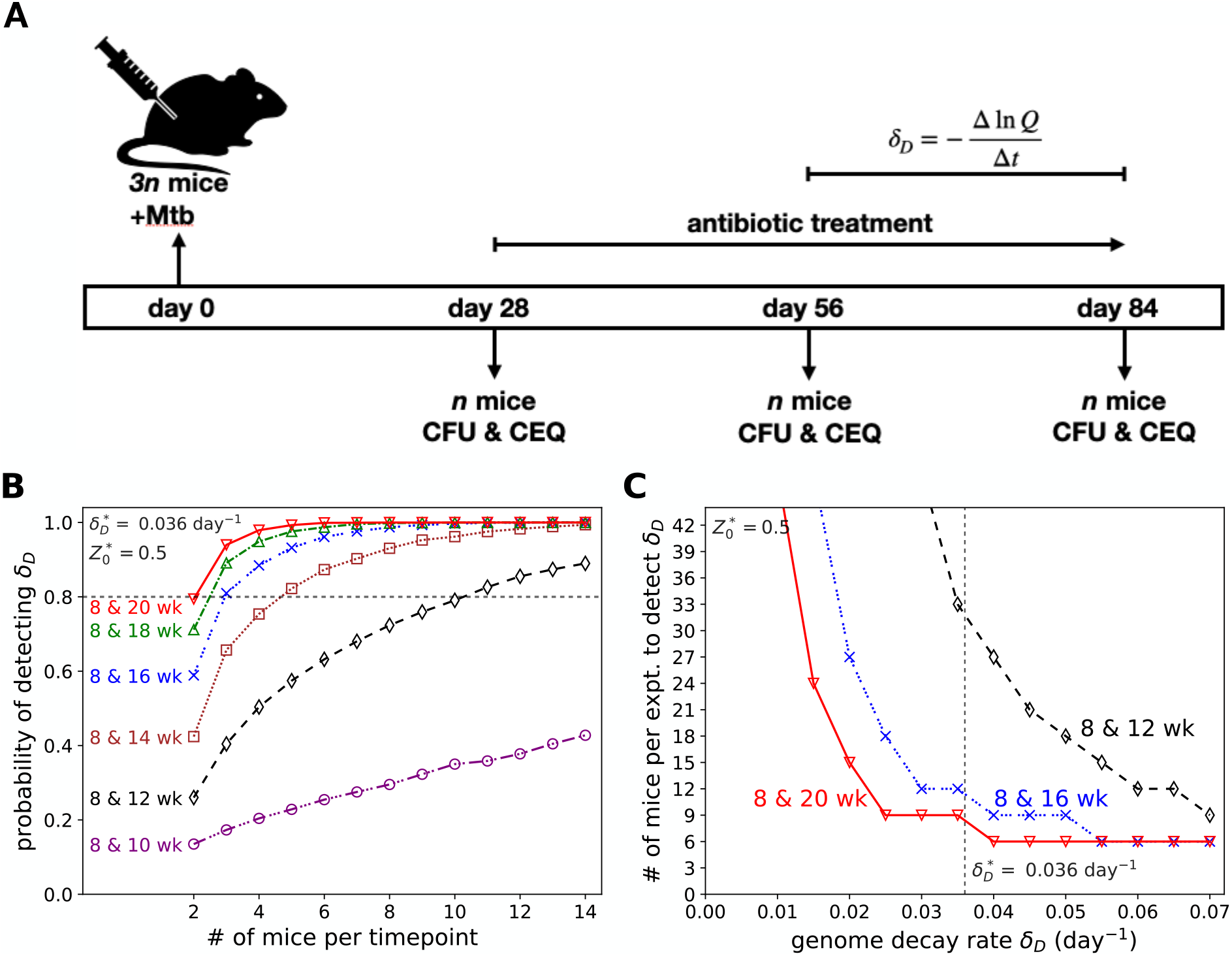
Power analysis for experiments to rigorously determine the decay rate of Mtb genomes *δ*_*D*_. **A**: Experimental design for measuring the decay rate of detectable Mtb genomes in lungs of infected mice. Mice (3*n*) are infected at day 0 and at day 28 (4 wks) post-infection, CFU and CEQs are measured in *n* mice while 2*n* mice start Ab treatment. At 56 days post-infection, CFUs and CEQs are measured in *n* mice. Finally, at some later time (e.g., 84 days post infection), CFUs and CEQs in the final *n* mice are measured. Day 56 and day 84 (or later) are then used to estimate the rate of CEQ decay if the number of viable bacteria at these time points is sufficiently small. **B**: Statistical power for detecting a statistically significant (*p <* 0.05) genome decay rate if *δ*_*D*_ = 0.036 day^−1^. We simulate experimental design in **A** with measurements of CFU and CEQs taken at day 56 (week 8) and other times (e.g., 10, 12, 14, 16, 18, or 20 wks post infection) with DD model and assuming log-normally distributed noise in measuring CFUs and CEQs estimated from Muñoz-Elías *et al*. ^6^ (see Materials and methods for more detail). In simulations we assume that ratio *Z*(28) = 0.5 at day 28 post infection (see **Supplemental Figures S3 and S4** for power analyses with other values of *Z*(28)). Horizontal line denotes power of 80%. **C**: The number of mice needed for 80% statistical power to detect different values of the Mtb genome decay rate *δ*_*D*_. Vertical dashed line denotes the value *δ*_*D*_ = 0.036*/*day (**Supplemental Figure S2**).

To evaluate the number of mice needed to estimate the Mtb genome decay rate of a particular value, we simulated Mtb dynamics in accord with the DD model using elements of the previously published data^6^. Specifically, we used the averages and standard deviations of the CFUs and CEQs at 28, 56, and 84 days post-infection from Muñoz-Elías *et al*. ^6^. We noted, however, that the mean value of CEQ numbers (*Q*) at 28 days post-infection was actually lower than the mean CFU value (*B*) – in the DD model this corresponds to negative amounts of killed bacteria. To correct for this discrepancy, we performed the analysis with different assumed CFU/CEQ ratios of 0.5 (**Figure 4**), 0.9 or 0.1 (**Supplemental Figures S3 and S4**, respectively) at 28 days post-infection.

Our results suggest that having *n* = 4 mice per time point (12 mice per experiment) is insufficient to accurately estimate the genome decay rate of *δ*_*D*_ = 0.036*/*day that is consistent with the reported result^6^. For the CFU/CEQ ratio *Z* = 0.5 at the start of treatment and measurements separated by 28 days (as in Muñoz-Elías *et al*. ^6^), we estimate that *n* = 11 mice per time point (or 33 mice per experiment) are needed detect genome decay rate equal to 3.6% per day (“8 & 12 wk” in **Figure 4**B). The number of mice needed to detect this decay rate drops substantially if the experiment is lengthened, with only *n* = 5 mice needed per time point (15 mice per experiment), if the experiment is extended by an extra two weeks (“8 & 14 wk” in **Figure 4**B). Results for other assumed CFU/CEQ ratios at start of treatment are conceptually similar, though the exact number of mice per experiment is affected by different shapes of the initial distribution of CEQs and by different influence of the viable population on the dynamics of CEQs (**Supplemental Figures S3 and S4**). Unsurprisingly, the number of mice needed to detect smaller genome decay rates begins to grow quite rapidly with decreasing decay rate (**Figure 4**C). Thus, to rigorously evaluate the Mtb genome decay rates with a relatively small numbers of mice there is a need to follow CEQ decay for longer times.

One may argue that measuring CFU and CEQ numbers at three different time points (at start of treatment, at intermediate time point, and at the end of treatment) is superfluous and measuring these numbers only at start and end of treatment may suffice to estimate the CEQ decay rate. Therefore, we performed another set of simulations where Mtb genome decay rate is detected by only using measurements at 28 and 84 days post-infection. Indeed, such experimental design would require fewer mice to achieve the same statistical power to detect a particular genome decay rate due to the longer time interval between measurements (**Supplemental Figures S5– S7**). However, because of the expected rise in CEQ numbers early after treatment start, our analysis suggests that such experimental design will result in bias toward underestimating the value of *δ*_*D*_, due to the dependence of the dynamics of *Q* on *B* (**Supplemental Figures S5–S7**). It is also possible that the viable population is underestimated in animals undergoing antibiotic treatment^20^, and precise degree of bias is difficult to estimate, since the genome decay rate is not precisely known. Furthermore, the possible presence of VBNC bacteria shortly after the start of antibiotic treatment may influence the apparent decay of Mtb genomes^21,22^. These effects will most likely compound the bias toward underestimating the decay rate of Mtb genomes from only two time points (start and end of treatment). For these reasons, experiments that allow to measure CFU and CEQ numbers at three time points (e.g., **Figure 4**A) would be more robust although at a higher number of mice.

### ID and DD models predict different dynamics at small infection doses

Our results so far suggested that both ID and DD models could accurately describe the data on Mtb dynamics in mice (**Figure 2**) or monkeys (**Figure 3**) but predict quite different rates of Mtb replication and death (**Tables 1 and 2**). These results, however, were based on deterministic predictions of the alternative models. It is well known that dynamics of Mtb is characterized by substantial variability. This is seen both among hosts^23–25^ and among lesions within individual hosts^13,26,27^. In studies with NHPs, differences in CFUs or CFU/CEQ ratio between individual lesions has been used to identify potential immune correlates of protection^5,13^. Unfortunately, differences in these quantities can potentially be influenced by a variety of causes, including different animals’ immune responses^28^, host and pathogen genetic variation^29^, timing of lesion formation^1,13,27^, timing of measurement^13,30^, and stochastic nature of replication and death processes. Disentangling stochastic noise from truly heterogeneous dynamics is challenging^31^. When lung lesions originate from just one or two bacteria, as is thought to be the case for individual granulomas^13,27^, a substantial contribution to variability in the trajectories results from the stochastic nature of replication and death, which itself is inevitable, given the stochastic nature of chemical reactions^25,31–33^. Therefore, we investigated whether simulating Mtb dynamics stochastically, assuming that infection starts with a single bacterium, results in different predictions by the ID and DD models.

To estimate the influence of stochasticity on dynamics of CFUs and CEQs in the alternative models, we performed stochastic (Gillespie) simulations of ID and DD models, using parameters derived from our fits to the data from individual lesions of macaques infected with a low dose of Mtb (**Figure 3** and **Table 2**). Specifically, we used the rates estimated with *δ*_*D*_ = *δ*_*Q*_ = 0.036 day^−1^, and with *Z* at 21 days post-infection equal to *Z* at 28 days post-infection (boldface rows in **Table 2**).

A few observations from the simulation results are noteworthy (**Figure 5**). Both models predict some variability in CFUs and CEQs due to the stochastic dynamics of replication and death (**Figure 5**Ai – Bii). Critically, the ID model predicts some lesions with CFU/CEQ ratio above one (*Z >* 1, **Figure 5**Ci), and in about 8% of trajectories, the CEQs decay to zero, while the CFUs survive, so that the CFU/CEQ ratio tends to infinity. (In visualizing the simulation results, we replaced values of 0 CEQs with 0.1, in order to allow visualization of CEQs on the log scale, and to allow visualization of diverging CFUs/CEQs on a finite scale.) Extremely large values of the CFU/CEQ ratio are biologically unreasonable even though CFU/CEQ ratios slightly higher than one have been observed experimentally^2,6,13,34^. This experimental observation could be understood simply as the result of some experimental noise; in this case, the ID model may be regarded as capturing some of the uncertainty inherent in experimental measurements. On the other hand, the observation of some lesions with CFU/CEQ ratio above one may actually point to a greater challenge, such as a systematic under-counting of Mtb genomes.

**Figure 5:**
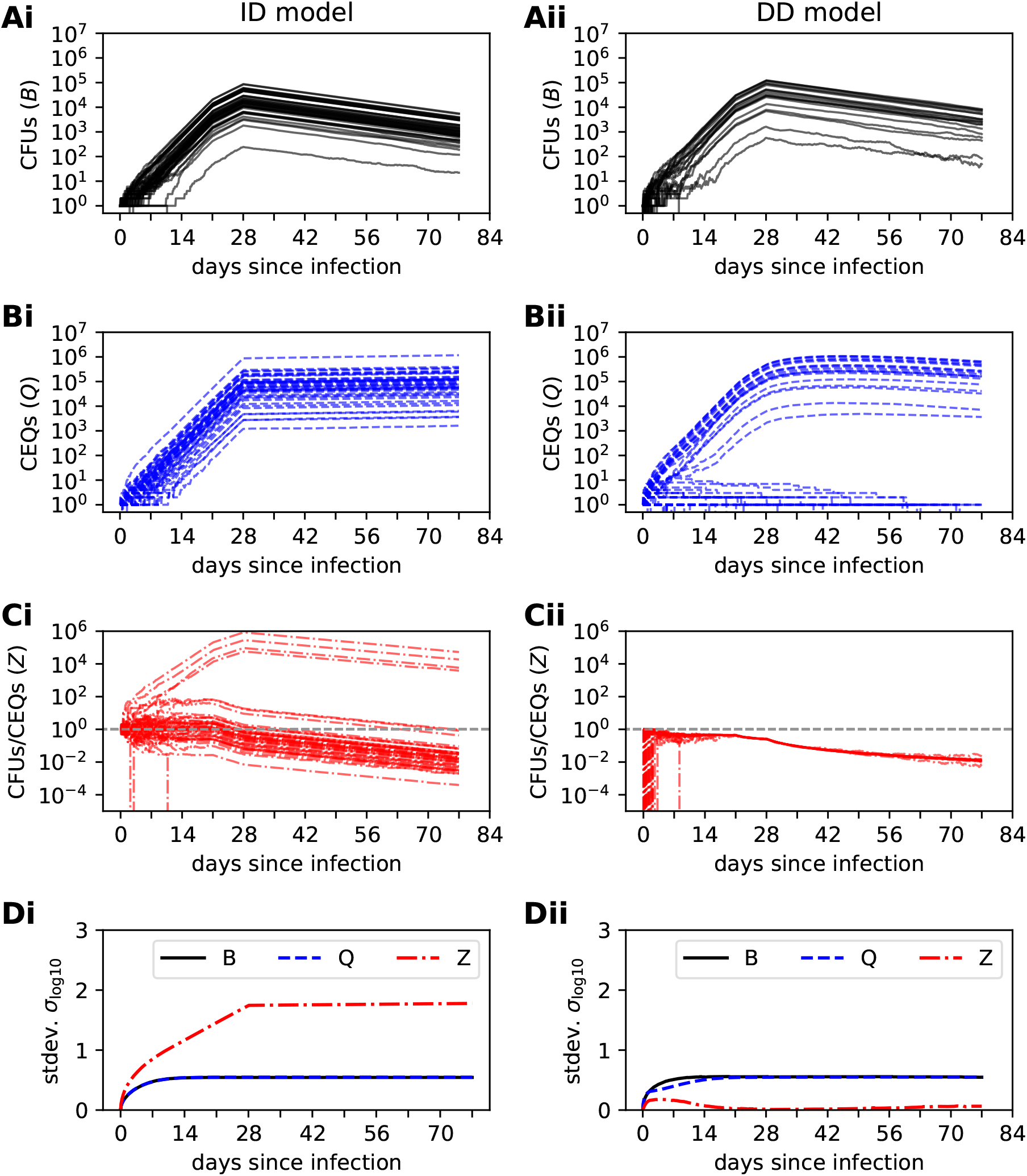
ID and DD models generate different predictions at small infection doses. We performed stochastic (Gillespie) simulations of ID (**Ai** - **Di, eqns. (6)–(7)**) and DD (**Aii** - **Dii, eqns. (8)–(10)**) models, using rates estimated by fitting the models to the Mtb CFU and CEQ values found in individual lesions in the lungs of macaques (see **Figure 3** and **Table 2**). We show the dynamics of the total number of bacteria *B* per lesion/granuloma (**A**), total CEQ number *Q* per lesion (**B**), CFU/CEQ ratio *Z* per lesion (**C**), and standard deviation of predictions for *B, Q*, and *Z*. In (C), the grey dashed line indicates *Z* = 1. In (A), (B), and (C), for visual clarity, we show results of only 50 simulations run for 77 days (11 weeks) each starting with *B*(0) = 1 and *Q*(0) = 1 (for ID model) and *B*(0) = 1 and *D*(0) = 0 (for DD model); other parameters are given in **Table 2** in rows for *δ*_*D*_ = 0.036*/*day (DD model) or *δ*_*Q*_ = 0.036*/*day (ID model) with “high early rates”. For accumulation of statistics in (D), we simulated 10,000 trajectories for each model.

While both models predict some variability in *B* and *Q* as a result of stochastic replication and death, the ID and DD models predict different variability in CFU/CEQ ratio *Z* over time. The ID model predicts substantially more variability in *Z* than in *B* and *Q* (**Figure 5**Di). On the contrary, after some initial stochastic noise produced when populations are small (first ∼ 5 days of infection), the DD model predicts narrowing of the distribution of *Z* over time (**Figure 5**Cii&Dii), suggesting that while stochastic dynamics account for a substantial portion of variability in CFUs and CEQs, variability in the CFU/CEQ ratio is primarily due to dynamical differences between lesions, and is minimally affected by stochastic nature of replication and death.

The latter result can be explained given the properties of the DD model; in the DD model the dynamics of the CFU/CEQ ratio *Z* approaches limiting behavior over time depending on whether the population is growing or declining. When bacterial numbers are increasing or are static (or declining more slowly than the genome decay rate), *Z* approaches the limiting value, *Z*_∞_ (**eqn. (S.10)** and **Figure 1**Fii). When bacterial burdens are decreasing faster than the genome decay rate, dynamics of *Z* approach exponential decay with rate *ρ* = *r* − *δ* + *δ*_*D*_ (**eqn. (S.14)** and **Supplemental Figure S1**). To better understand the convergent dynamics of *Z*, we calculated the time for log *Z* to decay halfway from its initial value (in the DD model, *Z*(0) = 1) to its asymptotic limit. Since it is typical to consider *Z* on a logarithmic scale, this time approximately reflects a “half life” of deviation of *Z* from limiting behavior, though it should be noted that variability in *Z* will not necessarily decrease exponentially. We performed this analysis with the DD model, with *δ*_*D*_ = 0.036 day^−1^.

We considered rates consistent with the dynamics of CFUs in two cases. First, for a case with increasing CFUs, we considered the dynamics of CFUs in B6 mice (dataset 2), between 4 and 8 weeks post-infection (**Supplemental Figure S8**A). Second, for a case with decreasing CFUs, we considered the dynamics of CFUs in macaques (dataset 3) between 4 and 11 weeks post-infection (**Supplemental Figure S8**B). In both cases, *r* and *δ* were varied in a feasible range, with *r* − *δ* fixed to agree with the net rate of population change (*i*.*e*., *r* − *δ* = Δ ln *B/*Δ*t*). In the former case, with the bacterial burden increasing, we find that the deviation of CFUs/CEQs from the asymptotic limit has a half life less than approximately two weeks. And for the macaques, with decreasing bacterial burden, we find a half life equal to or less than about 3 weeks for the whole range of possible rates, and a half life of about 1 week in the range of our best estimate of *r* and *δ*.

When *B* is increasing (**Supplemental Figure S8**A), the result reflects the approximate memory length of variability in *Z*. On the contrary, when rates are decreasing (**Supplemental Figure S8**B), it reflects the approximate timescale over which dynamics will be expected to converge to the behavior of the ID model (exponential decay of *Z*). Note, however, that since the exponential decay will still have an intercept that depends on initial *Z*_0_ (**eqn. (S.14)**), variability in *Z* is expected to be preserved when CFUs are decreasing, even after convergence to exponential decay. Taken together, these results lead to the prediction that when average bacterial burdens are increasing or static in an experiment, variability in *Z*, quantified, for example, as the variance of log *Z*, will tend to decrease, and when average bacteria populations decline in an experiment, variability in *Z* will tend to be static or increase over time. If experiments suggest opposite dynamics (e.g., increase in variance of log *Z* when CFUs are increasing or constant), it would reject this version of the DD model assuming that parameters for replication and death are identical between individual granulomas.

## 4 Discussion

Studies of mice^6^, monkeys^13^, and rabbits^1,2^ have utilized measurement of CFUs and CEQs to probe in-host dynamics of Mtb. While Mtb dynamics vary among different animal models, studies utilizing CFUs and CEQs of Mtb have concluded that after the onset of host adaptive immunity, the majority of the Mtb population is in a non-replicating or dead state^8^. In addition to using different mathematical models, by assuming that Mtb genomes do not appreciably decay, studies in mice and monkeys have generally treated CEQs as a surrogate for CBB. No consistent rigorous modeling framework for analyzing data, based on measurements of CFUs and CEQs, has been developed.

To address this research gap, we developed a general and two simplified alternative models of the in-host dynamics of CFUs and CEQs of Mtb (**Figure 1**); the general model includes several important processes of the CFU and CEQ dynamics but it was overparameterized and was not suited to be rigorously fit to experimental data. The two extreme cases of the general model make different assumptions of how CEQs are generated during the infection: the independent dynamics model assumes that CFUs and CEQs replicate and die independently (**eqns. (6)–(7)**) while the dependent dynamics model assumes that CEQs in excess of CFUs originate exclusively from dying bacteria (**eqns. (8)–(10)**). By using data from Mtb-infected mice treated with INH we estimated that Mtb genomes are not immortal but decay at an appreciable rate of *δ*_*Q*_ = *δ*_*D*_ = 0.036*/*day (**Supplemental Figure S2**) which is similar to a value previously reported for Mtb in granulomas of macaques^13^. Importantly, independently of the value of Mtb genome decay rate, both ID and DD models could accurately describe the CFU and CEQ data in mice (**Figure 2**) or monkeys (**Figure 3**) suggesting that these data alone are insufficient to discriminate between the alternatives.

Given the estimated Mtb genome decay rate of *δ*_*Q*_ = *δ*_*D*_ = 0.036*/*day, we found that Mtb replicates and dies at substantial rates during chronic infection (4-16 weeks) in mice (**Figure 2** and **Table 1**) or acute infection (first 3 weeks) in monkeys (**Figure 3** and **Table 2**). Simulating Mtb dynamics stochastically (assuming that infection starts with one bacterium) resulted in different predictions between ID and DD models. Specifically, the ID model predicted a large range of the CFU/CEQ ratios with about 8% of simulations predicting *Z* » 1 which is biologically implausible (**Figure 5**). In contrast, in the DD model, while CFUs and CEQs exhibited highly stochastic dynamics, the CFU/CEQ ratio was highly constrained (**Figure 5**); this can be explained by asymptotic behavior of the model (**eqn. (S.10)** and **eqn. (S.13)**). Finally, by using the DD model we performed power analysis that predicted sampling times and the number of mice needed accurately to estimate the Mtb genome decay rate of a particular value (**Figure 4**).

The models we developed here have some attributes in common with existing models. The ID model is conceptually similar to the Lin *et al*. ^13^ model, but with explicit separation of dynamics into contributions from replication and death, and with explicit inclusion of the genome decay rate *δ*_*Q*_. Our analysis of these models revealed difficulties with connecting model parameters to physical processes, so that fitting with the ID model or Lin *et al*. ^13^ model requires care in interpretation of results. Likewise, the DD model is conceptually similar to the model of Muñoz-Elías *et al*. ^6^, but includes decay of genomes *δ*_*D*_ and is formulated with ODEs. Inclusion of genome decay in the model revealed strong sensitivity of estimated Mtb replication and death rates to the assumed rate of genome decay.

Our estimate of 3.6% per day for the decay rate of detectable Mtb genomes in lungs of B6 mice is close to the approximately 4% per day estimated previously for lesions from macaques’ lungs, using time matched samples pre- and post-treatment with INH^13^. Interestingly, viable Mtb has a similarly slow decay rate in soil (decay from 10^8^ CFU/g to 2 × 10^3^ CFU/g in 12 months resulting in the decay rate *d* = 3%*/*day, ref^35^; see also Supplemental Information for a slightly more sophisticated analysis of Mtb decay in soil). Our power analysis suggests that experiments in which CEQs are measured only at two time points (at start and end of treatment) would provide biased estimates of the Mtb genome rate (**Supplemental Figure S5**) suggesting that 4% per day genome decay rates found in macaques may be an underestimate. In view of the non-zero Mtb genome decay rate, caution should be exercised in calibrating models using CEQ data under the assumption that Mtb genomes are immortal^3^.

Our estimates of the replication and death rates of Mtb in chronically infected (4-16 weeks) mice (**Table 1**) are slightly smaller than, but similar to, those predicted by experiments using a replication clock plasmid^10,12^. To quantify this further, we used the rates reported by McDaniel *et al*. ^12^ to fit the DD model to the CEQ and CFU data for B6 mice from the Muñoz-Elías *et al*. ^6^ study. Under this constraint, the best fit of the DD model to dataset 1 was achieved when *δ*_*D*_ ≈ 0.07 day^−1^, the upper bound of the 95% confidence interval we estimated for *δ*_*D*_ (not shown). These results thus reconcile different interpretation of the rate of Mtb replication and death in chronically infected mice and obtained using CEQs or replication clock plasmid^6,10,12^.

When working with multiple models, it is useful to consider evidence and arguments that may help to discriminate between alternatives. The convergence of different trajectories toward a single limiting value of CFUs/CEQs as predicted by the DD model in stochastic simulations (**Figure 5**Cii), is notably not seen in experiments, e.g., there is large variability in the CFU/CEQ ratio in individual granulomas of monkeys throughout the experiment. This observation alone makes it clear that the DD model assuming identical parameters for each granuloma, in which variability in CFUs and CEQs arises only via stochastic dynamics, is inconsistent with such data.

In contrast, change in the variance of log of CFU/CEQ ratio in granulomas of rabbits appears to follow predictions of the DD model, e.g., in rabbits variance of the CFU/CEQ ratio decreased when average bacterial burdens increased and increasing as bacterial burdens decreased (*e*.*g*., see Figure 1A&E of Blanc *et al*. ^1^). Extending the DD model to allow for subpopulations of viable bacteria with different replication kinetics may allow to make the DD model match the data more accurately. The ID model, on the other hand, despite difficulty in associating it with a simple biological mechanism, captures more of the variation observed in experiments, including occasional observation of CFUs/CEQs *>* 1. Even the prediction of some samples with CEQs = 0 while CFU numbers change, though it seems biologically unreasonable, could be viewed as reflecting the high detection limit of CEQs^36^, compared with CFUs. The FID model (**eqns. (11)–(12)**) may be useful for describing dynamics of CFUs and CEQs in a situation with CFUs « CEQs. When CFUs and CEQs are separated by multiple orders of magnitude, the dynamics of CEQs are expected not to be significantly influenced by changes in CFUs, which would contribute only a very small fraction of the total population of genomes. In this case, the CEQs and CFUs really are expected to evolve independently of one another, and modeling them in that manner is probably appropriate. Note, however, that the ID model in the strict form (equal replication rates for the two populations) is not quite appropriate for this purpose, since it requires CEQs to replicate at the same rate as CFUs. To avoid the extra parameter in the FID model, the model could be constrained differently, for example, by fixing the replication rate of CEQs to zero, as was done by Lin *et al*. ^13^ for the second stage of their model (4 weeks post-infection and later). Note, however, that the DD model automatically produces this limit, but without introducing an extra parameter.

Our work has several limitations. While we have considered several mathematical models, the formalism considered here does not encompass the full range of possible dynamics of CFUs and CEQs in animals. The two models emphasized in this work represent extremes that are related to the most common ways in which CFUs and CEQs have been discussed in the literature (especially Lin *et al*. ^13^ and Muñoz-Elías *et al*. ^6^), and do not account for the likely case of heterogeneous Mtb dynamics^16,37,38^. At one extreme, the DD model considers all bacteria to be either viable and culturable or dead. On the other hand, the ID model treats all viable bacteria as replicating with the same rate, even if they are not culturable (they share a single replication rate). This leaves out the likely scenario involving VBNC bacteria, or other “dormant” bacteria that persist in a mostly non-replicating state; the reality is almost certainly between these extremes. Indeed, McDaniel *et al*. ^12^ found that data from replication clock experiments with mice could be explained by a substantial fraction (up to 25% at day 111) of non-replicating, but still culturable, bacteria, with most bacteria replicating at high rates; notably, this fraction of non-replicating bacteria could be much higher, if they are also non-culturable (since they will not be detected at all in such experiments)^12^. However, the formalism developed here in the DD model can be adapted easily for a heterogeneous population, though at the expense of a significant increase in the number of model parameters.

It is also possible that the decay rate of detectable genomes varies between different tissue types (or even between stages of infection). For example, in control (untreated) Mtb-infected rabbits median CEQs in uninvolved lung tissue dropped by more than two logs_10_ between 12 and 16 weeks following Mtb infection, while median CEQs in cellular lesions dropped by less than one log_10_ over the same time period^2^. However, because CEQs were not detected in a large portion of lesions, CEQ decay rate may not be reflected in median CEQ values.

Our approach here does not address some possible factors that could influence production, decay, and measurement of CEQs. As pointed out in Muñoz-Elías *et al*. ^6^, there is a possibility of destruction of genomes concurrent with bacteria killing, as one might expect, for example, as a result of phagocytosis. While our general model includes this as a possibility, it is not featured in the simpler models: in the ID model, it is only included implicitly, rolled into the parameters, and it is excluded in the DD model. This may lead to fewer CEQs from dead bacteria than expected, when a substantial amount of killing occurs, which may explain some of the discrepancy between the rates estimated here for chronically infected B6 mice and the estimates using a replication clock plasmid^10,12^.

It is also possible that detection limit of CEQs depends on the state the bacterial chromosome is in, for example, from being cell-free DNA molecules in a tissue to DNA molecule inside of dead bacteria with either intact or damaged cell walls, or within VNBC bacteria. As Mycobacteria are known to be challenging to lyse, such heterogeneity may lead to differences in detectability of genomes, in addition to heterogeneous decay of chromosomes^39^. Although noise must also play some role, this might help to explain occasional measurement of CFUs greater than CEQs in lesions from macaques^13^, lesions from rabbits^2^, whole lungs from mice^6^, and even several successive average values from *in vitro* experiments with *Mycobacterium abscessus* ^34^. This is also a possible explanation for the large values of *r* and *δ* needed to fit the data between 4 and 8 weeks postinfection in B6 mice treated with antibiotics (**Supplemental Figure S2**). Furthermore, the relatively high detection limit of CEQs^36^, around 10^3^ per granuloma, could skew statistics toward higher estimates of mean CEQs. Since estimated rates depend on small differences in CEQs (e.g., **eqns. (S.18)–(S.19)**) between different time points, such potential sources of bias may have significant effects on estimated rates.

The framework used here has only been applied to averaged data and published error bars. While we consider this a necessary refinement of the existing models of average dynamics, further comparison of model predictions with data from individual animals (e.g., individual mice or granulomas of monkeys/rabbits) would be very useful We anticipate that this will require extension of the models to include heterogeneous bacteria populations (with, for example, subpopulations of viable bacteria, or a continuous distribution of replication and death rates)

Because previous studies in mice and macaques did not quantify CEQs prior to 4 weeks postinfection^6,13^, it is difficult to accurately estimate replication and death during this time period. We have addressed this by exploring the sensitivity of rates to changes in assumed values of CEQs at 2 weeks (mice) or 3 weeks (macaques) post-infection (**Tables 1 and 2**). Nevertheless, measurement of CEQs at early times after infection would be valuable for more precise quantification of acute-phase replication and death rates.

Our work opens avenues for future research. While only mean and median CFUs and CEQs were considered in the present work (due to lack of data availability for individual animals), additional information on heterogeneity of population dynamics may be gained by considering whole distributions among animals and/or lesions from individual animals. Data from pharmacological studies in rabbits may help to shed light what can be learned from considering the whole distribution of CFUs and CEQs among different lesions and different animals; large datasets have been published including individual lesions from rabbits treated with different antibiotic regimens^1,2^. Furthermore, application of modeling to these datasets may help to quantify modes of action of antibiotic treatments. Ultimately, it may be useful to extend this work to analysis of biomarkers from human patients, in a clinical setting. Detection of Mtb chromosomes in sputum as a marker of antibiotic efficacy has been previously explored^40^. Consistent with the observations of Muñoz-Elías *et al*. ^6^ and with the prediction of the DD model (**Supplemental Figure S2**), a possibly counterintuitive increase in detectable genomes in sputum was observed after treatment start, followed by decrease of detected Mtb chromosomes over time^40^. This observation could also be explained by ongoing Mtb replication (as in the DD model), by a change in the detectability of genomes, or by a change in the number of detectable chromosomes per bacillus due to antibiotic-mediated block of cell growth.

While further studies are needed to falsify possible models, at this point, we cautiously find the DD model to be of greater practical use in most situations. Because it is more constrained and is simpler to interpret in terms of model biological processes, differences between observations and DD model predictions lead readily to a path forward with new hypotheses (in the present case, for example, modeling subpopulations of viable bacteria). This advantage notwithstanding, an analysis similar in style to our FID model or the Lin *et al*. ^13^ model is useful for describing overall dynamics, as long as one maintains appropriate awareness of the difficulties of interpreting such a model in terms of fundamental biological processes.

While CEQs are challenging to measure due to the presence of just one chromosome per bacterium, messenger (**mRNA**) or ribosomal RNA (**rRNA**) is also quantifiable by qPCR and is present in significantly larger numbers in bacterial cells^38,41,42^. Counts of Mtb RNA has been suggested as a robust surrogate for CFUs in preclinical evaluation of treatments and for evaluation of treatment endpoints in clinical applications, since RNA from VBNC bacteria can be detected^40,41^. The DD model could be extended for CFUs and rRNA dynamics^43^, with the dynamics of total rRNA counts determined by summing viable and dead contributions, and introducing a factor or factors to account for the number of rRNA counts per cell, which may vary over time, *e*.*g*., as a function of replication rate^44^. Application of such a modified DD model may help in applying rRNA counts to estimate the probability of relapse following long-term treatment when CFU numbers reach zero.

Given the different underlying mechanisms and the different estimated rates from the ID and DD models, the fact that both ID and DD models can fit data for mice and for macaques indicates that CFUs and CEQs alone are not sufficient to fully define the in-host replication state of bacteria. At a conceptual level, this is inevitable: while the present work aims to clarify evaluation of the in-host dynamics for simplest-case models, there is substantial experimental evidence of significantly more complexity in real systems ^2,16,38^. Studies are needed that evaluate the dynamics of CFUs and CEQs with the aid of models that include multiple subpopulations with different replication and/or death kinetics; the DD model could be adapted for this purpose. However, inclusion of multiple subpopulations rapidly increases the number of fitted parameters needed to evaluate model dynamics. For this reason, pairing of CFU and CEQ measurements with additional experimental probes of bacteria replication and/or death, e.g., replication clock plasmid, may help in characterizing in-host dynamics of Mtb more accurately.

Various possibilities exist for providing additional information. For example, fluorescently tagged replisome components can provide spatial information about heterogeneous replication^45^, and DNA sequence read coverage can be connected with the rate of replication using mathematical modeling^46–48^. The ratio of short-lived pre-ribosomal RNA to longer-lived rRNA (RS ratio)^38^ appears to be related to replication rates of Mtb by indicating ongoing ribosome synthesis, though mathematical models are still needed to rigorously connect the RS ratio to replication rates. Use of a replication clock plasmid^10^ along with CFUs and CEQs would likely aid substantially in quantifying replication and death dynamics with greater clarity. When the replication clock plasmid is used, a declining percentage of plasmid-containing cells over time indicates ongoing replication, while dynamics of CEQs and CFUs, in principle, indicates both replication and death. Simultaneous use of both techniques would provide the additional information needed to discriminate between models, allowing greater clarity in evaluation of in-host dynamics of Mtb, for development of improved vaccines and treatments for Mtb infection and TB.

## Abbreviations

CFU: colony forming unit
CEQ: chromosomal equivalent
CBB: cumulative bacterial burden
DD: dependent dynamics
ID: independent dynamics
GM: general model
FID: exible independent dynamics
Mtb: *Mycobacterium tuberculosis*
TB: tuberculosis
rRNA: ribosomal RNA
pre-rRNA: pre-ribosomal RNA
RS ratio: ratio of pre-rRNA to rRNA
INH: isoniazid
VBNC: viable but not culturable
ODE: ordinary differential equation
mSSA: stochastic simulation algorithm
qPCR: quantitative polymerase chain reaction
NHP: nonhuman primate

## Data sources

The data from the paper (digitized from original publications) along with the codes are available on github: https://github.com/allanfriesen/mtbCfuCeqDynamics.

## Code sources

All analyses have been primarily performed in python (ver 3.11). Example codes are available on the github (see link above).

## Ethics statement

No animal work has been performed.

## Author contributions

ADF and VVG conceived the overall concept of the study and developed alternative models. ADF performed all major analyses of the models including fitting models to data and stochastic simulations. ADF wrote the first draft of the paper and all authors read, edited, and agreed on the final version.

## Acknowledgments

We contacted lead authors of Muñoz-Elías *et al*. ^6^ paper for access to primary data from their study; however, the data could not be located. We do appreciate that the authors attempted to find the data.

## Supplemental Information

### S1 Additional mathematical considerations

#### S1.1 General Model

We developed a general model (**eqns. (3)–(5)**) of dynamics of CFUs and CEQs that includes the following components and processes (see **Figure 1**A and **eqns. (3)–(5)**):

- Subpopulations *B* and *D* represent culturable bacteria and unculturable bacteria, respectively. Unculturable subpopulation *D* includes both VBNC bacteria and chromosomes of dead bacteria.
- Culturable bacteria replicate with average per capita rate *r*.
- Culturable bacteria leave the culturable state with per capita rate 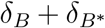, where *δ*_*B*_ is the conversion rate of *B* to dead and/or dormant state *D*, with (temporary) preservation of the genome, and 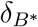 is the rate of bacteria death with simultaneous degradation of the genome (for example, by phagocytosis).
- Bacteria in state *D* may replicate with per capita rate *r*_*D*_, which cannot exceed *r*_*B*_.
- Genomes in subpopulation *D* are degraded with per capita rate *δ*_*D*_.

The dynamics of CFUs/CEQs = *Z* in the general model follow a logistical form:

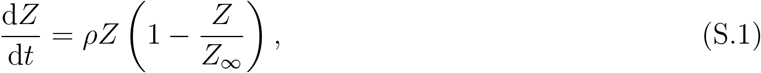

where

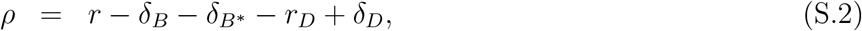

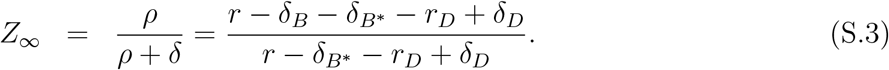

#### S1.2 Independent Dynamics model

##### S1.2.1 Interpretation of ID model parameters

Intermediate between the general model and the independent dynamics (**ID**) models we can define the flexible independent dynamics model (**FID, eqns. (11)–(12)**), obtained from the general model by setting 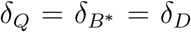 (**eqns. (12)–(11)**). In this model, the populations *B* and *Q* each have their own replication and death rates, resulting in 6 adjustable parameters including initial conditions. In order to fit experimental data unambiguously, an additional constraint is needed. The ID model is a special case of the FID model with the restriction that populations *B* and *Q* share the same replication rate, *r* = *r*_*B*_ = *r*_*Q*_. During early infection, this is a natural choice, since we expect *B* = *Q* at that point. It is possible that some other choices may make better sense later during infection. For example, the analysis of Lin *et al*. ^13^ appears to suggest that *r*_*Q*_ = *r* ≈ 0 after about 4 weeks of infection, although this is not entirely clear, since they do not list parameters.

In the ID model, *r* clearly represents the replication rate of viable bacteria; however, requiring that this constant also defines the replication rate of *Q*, the ID model in its restricted form implies that subpopulation *Q* is composed of a combination of platable and VBNC bacteria, all replicating with rate *r*. The rate *δ* cannot be thought of as representing a death rate in this model, since the underlying mechanism implies that the non-platable bacteria can replicate.

This difficulty is present even in the more flexible FID model. To see this, parameters of the FID and the DD models can be related to each other. The rate *r*_*Q*_ is simply the weighted average of *r*_*B*_ and *r*_*D*_:

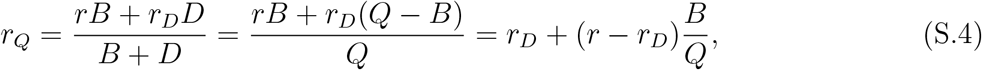

and *δ*_*Q*_ is conceptually similar to *δ*_*D*_, except that it represents an average decay rate of *all* genomes, rather than just those of dead bacteria:

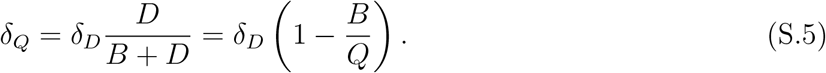

Because the difference *Q* − *B* changes over time, the parameters of both models cannot generally be taken simultaneously to be time-independent. The result of this analysis is that independent fitting of exponential (or logistical) functions to *B* and *Q* produces parameters that cannot be associated in a simple way with replication or death rates of particular populations.

#### S1.3 The Lin et al. model is not consistent with division of CEQs into platable and dead populations

The model of Lin *et al*. ^13^ (**eqns. (1)–(2)**) applies separate logistical forms for dynamics of CFUs and CEQs of Mtb in lesions from the lungs of macaques. Early during infection, the CFUs and CEQs are far smaller than their corresponding carrying capacities, so that the dynamics of this model is equivalent to that of the ID model. As such, it is also subject to the same difficulties with associating the rates with replication and death, as well as with the assumption that the VBNC population is negligibly small. However, if replication and death rates are allowed to change arbitrarily over time, one can derive their time dependence under the assumption that the CEQs divide cleanly into dead and platable bacteria, by application of **eqns. (S.24)–(S.25)** to the curves (**eqns. (1)–(2)**). The result is that both replication and death rates increase exponentially as *B* begins to saturate, then both rates decrease exponentially as *Q* saturates.

##### S1.3.1 Dynamics of CFUs/CEQs in FID and ID models

For the FID model, the dynamics of *Z* = CFUs/CEQs follow exponential decay at a rate that depends on replication and death rates of CFUs and CEQs:

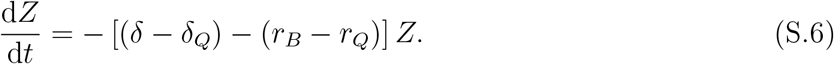

Because the rate of exponential decay depends on all four rates, association of the dynamics of *Z* with particular biological processes is difficult, without added assumptions. The ID model’s simplifying assumption that *r*_*B*_ = *r*_*Q*_ leads to the simpler result,

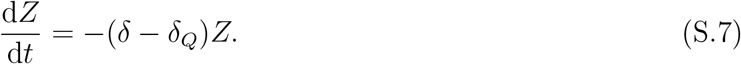

Under this assumption, decay of CFUs/CEQs reflects the difference between *δ* and the genome decay rate *δ*_*Q*_. If additionally, *δ*_*Q*_ were negligibly small, the dynamics of *Z* could be associated with the death/decay rate *δ* of CFUs.

#### S1.4 Dependent Dynamics model

##### S1.4.1 Interpretation of DD model parameters

Starting again from the general model, setting 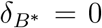 and *r*_*D*_ = 0 leads to the DD model, defined by **eqns. (8)–(10)**. Because the DD model contains explicit live and dead subpopulations in its formulation, interpretation of parameters in the DD model is simple: *r* represents the per capita replication rate of viable bacteria, *δ* represents the per capita death rate, and *δ*_*D*_ represents the per capita decay rate of detectable genomes of dead bacteria. It is also straightforward to allow some of the population *D* to replicate by introducing a small replication rate, *r*_*D*_ to the model. Additional data would be needed to justify addition of this parameter. *r*_*D*_ would, in principle, represent the average per capita replication rate of bacteria in subpopulation *D*; similarly, *δ*_*D*_ would now represent a degradation rate of genomes, averaged over both VBNC and dead populations. In such a case, the rates *r*_*D*_ and *δ*_*D*_ would be expected to vary over time, even when *r* and *δ* are constant, since the proportion of *D* that represents dead bacteria would most likely change over time. A model that explicitly distinguishes VBNC and dead populations would seem more realistic, though such a model likely would require additional fitting parameters, increasing the chance of overfitting.

##### S1.4.2 Dynamics of CFUs/CEQs in DD model

Similar to the general model, the dynamics of CFUs/CEQs = *Z* in the DD model follow a logistical equation:

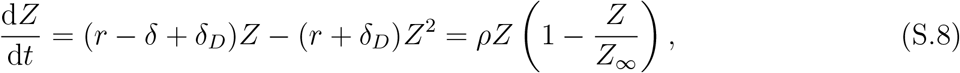

where net rate *ρ* and carrying capacity *Z*_∞_ are given by:

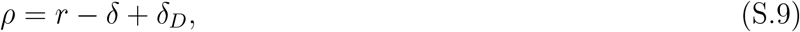

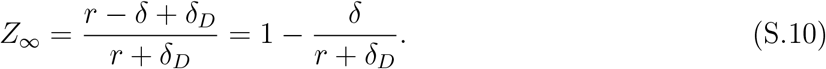

When bacterial numbers are increasing, *Z* saturates at *Z*_∞_. However, when bacterial numbers are decreasing, *Z*_∞_ is typically negative, so that the trajectory of *Z* instead approaches an exponential decay (**Supplemental Figure S1**).

During chronic infections, the net rate *ρ* is small, so that the first order term in **eqn. (S.8)** is small, and the long-time limit of *Z* is close to zero. In this case, the dynamics of *Z* are approximately second order.

Integration of the DD model gives the following forms for the dynamics of CFUs, *B*, dead bacteria genomes, *D*, and their ratio, *Z*:

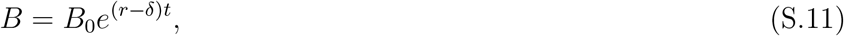

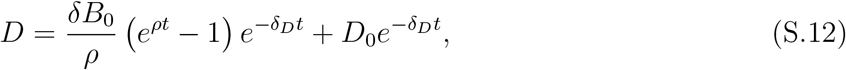

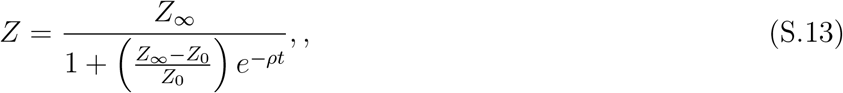

where *ρ* and *Z*_∞_ are defined as above.

When the viable population declines faster than the decay rate *δ*_*D*_ of detectable genomes of killed bacteria, the carrying capacity, *Z*_∞_, becomes negative. In this case, the long-time dynamics of *Z* asymptotically approach an exponential decay, described by the limiting behavior, *Z*_*L*_, of **eqn. (S.13)** when *t* is large and *ρ* is negative (**Supplemental Figure S1**):

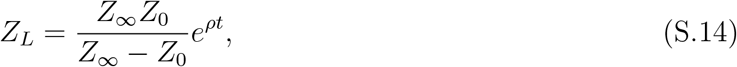

where as before, *ρ* = *r* − *δ* + *δ*_*D*_. When considered over a sufficiently long time, the full dynamics of decline of *Z* in the DD model will be approximated well by **eqn. (S.14)**. In this case, the difference in per capita rates of decline of *Z* and *B* provide an estimate of *δ*_*D*_:

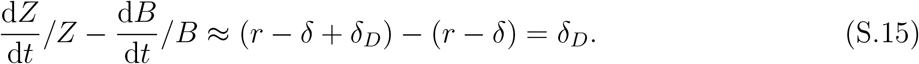

Use of this approximation slightly biases toward underestimate of *δ*_*D*_, so that it may be thought of as a method for estimating a *lower bound* on the degradation rate of Mtb genomes in vivo. However, this method only works if the dynamics of CFUs and CFU/CEQ ratio approaches an asymptote, e.g., for constant (time-independent) parameters; if the rate of Mtb replication and death change over time, this approximation may give incorrect estimate of the CEQ genome rate. As an example, we estimated the change in log *Z* = −0.104*/*day and log *B* = −0.056*/*day between 4 and 11 weeks in Mtb-infected macaques^13^; this gave a negative estimate of the genome decay rate *δ*_*D*_ = −0.104 − (−0.056) = −0.048*/*day highlighting the issue.

#### S1.5 Expressions for Estimating Model Parameters from Experimental Data

Ideally, rates would be extracted from time series including several different points. However, since studies usually include only two or three time points, it is more realistic to consider how parameters can be estimated from just a pair of points. In these analyses we assume that the CEQ decay rate is known.

For the ID model, *r* and *δ* can be estimated from the following expressions:

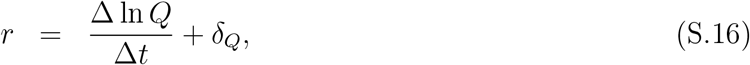

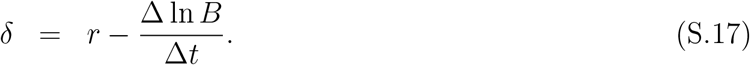

Likewise, for the DD model,

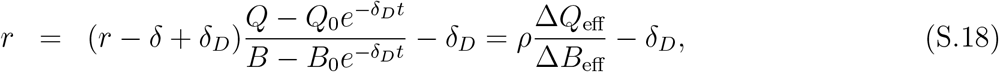

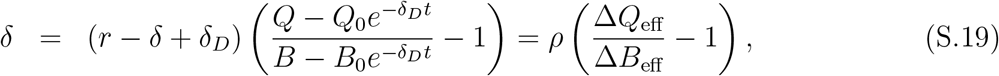

where

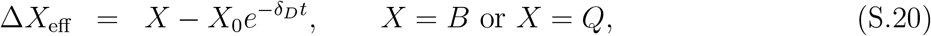

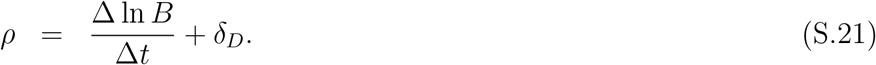

The following expressions could be used to produce the corresponding *r* and *δ* as a function of time for the for the ID model:

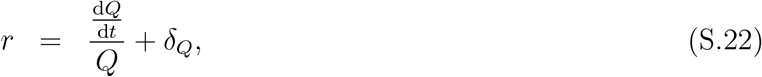

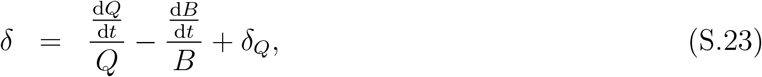

And for the DD model:

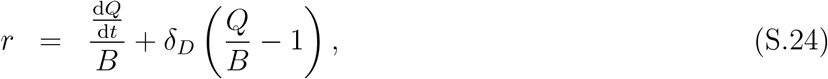

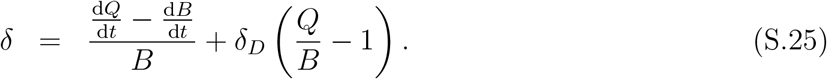

Note that while we have carried out this analysis for both ID and DD models, for completeness, the meaning of time-dependent *r* and *δ* in the ID model is unclear since *r* and *δ* are already aggregate variables, implying time-dependent replication and death rates.

### S2 Decay of viable Mtb in soil

We digitized measurements of Mtb CFUs isolated from soil samples at 1 month intervals for 12 months after inoculation with a defined number of Mtb bacteria (Figure 1 of Ghodbane *et al*. ^35^). We found that the viable population declines in two phases. During the first two months after inoculation (phase 1), CFUs/g declined with a half-life of about 4.9 days (decay rate *d*_1_ = 0.14*/*day). Subsequent decay in months 10-12 (phase 2) was substantially slower, with a half-life of about 36 days (decay rate *d*_2_ = 0.019*/*day). Assuming a single exponential decay and fitting both slope and intercept resulted in an estimated half-life of 22 days (decay rate *d* = 0.032*/*day) that is quite similar to our estimated half-life time of CEQs in mice. If, instead, the intercept was fixed to the initial value of 10^8^ CFUs/g, the resulting slope was steeper, corresponding to a half-life of about 14 days (decay rate *d* = 0.050*/*day).

## S3 Additional figures

**Supplemental Figure S1:**
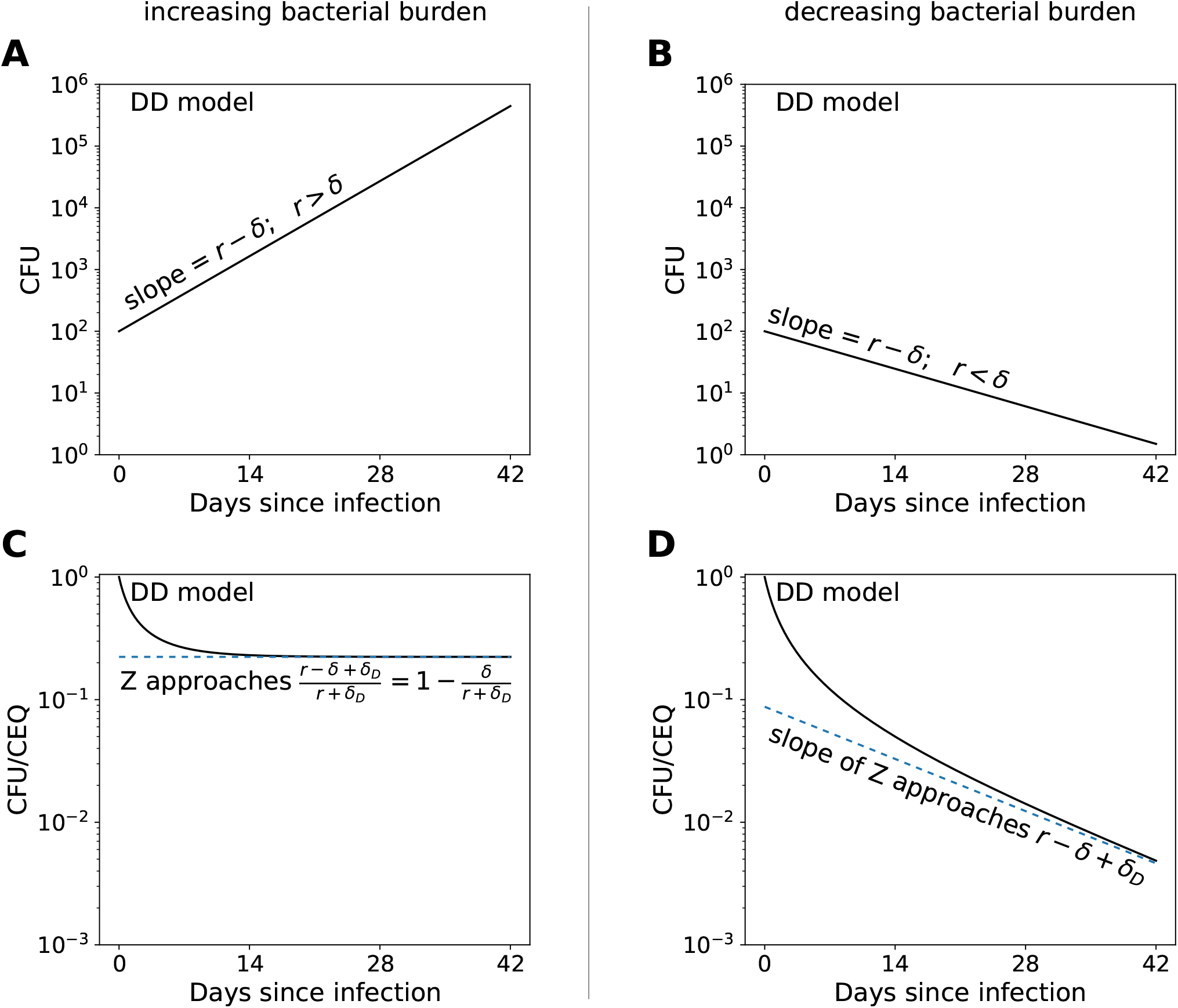
DD model predicts different asymptotic behavior for CFU/CEQ ratio *Z* for expanding vs. contracting bacterial populations. We show predictions of the DD model (**eqns. (8)–(10)**) when bacterial population grows in size (*r > δ*, panels **A** and **C**) or when it declines in size (*r < δ*, **D** and **D**). More precisely, when *r* + *δ*_*D*_ *> δ, Z* approaches a positive carrying capacity (**C**). When *r* + *δ*_*D*_ *< δ*, the dynamics of *Z* approach the asymptotic limit, *Z*_*L*_ = (1 − 1*/Z*_∞_))^−1^ exp(*ρt*) (**D**). In these simulations, parameters of the model are *r* = 1 day^−1^ (panels A and C), *r* = 0.7 day^−1^ (panels B and D), *δ* = 0.8 day^−1^, *δ*_*D*_ = 0.03 day^−1^, *B*_0_ = 10^3^, and *D*_0_ = 0.

**Supplemental Figure S2:**
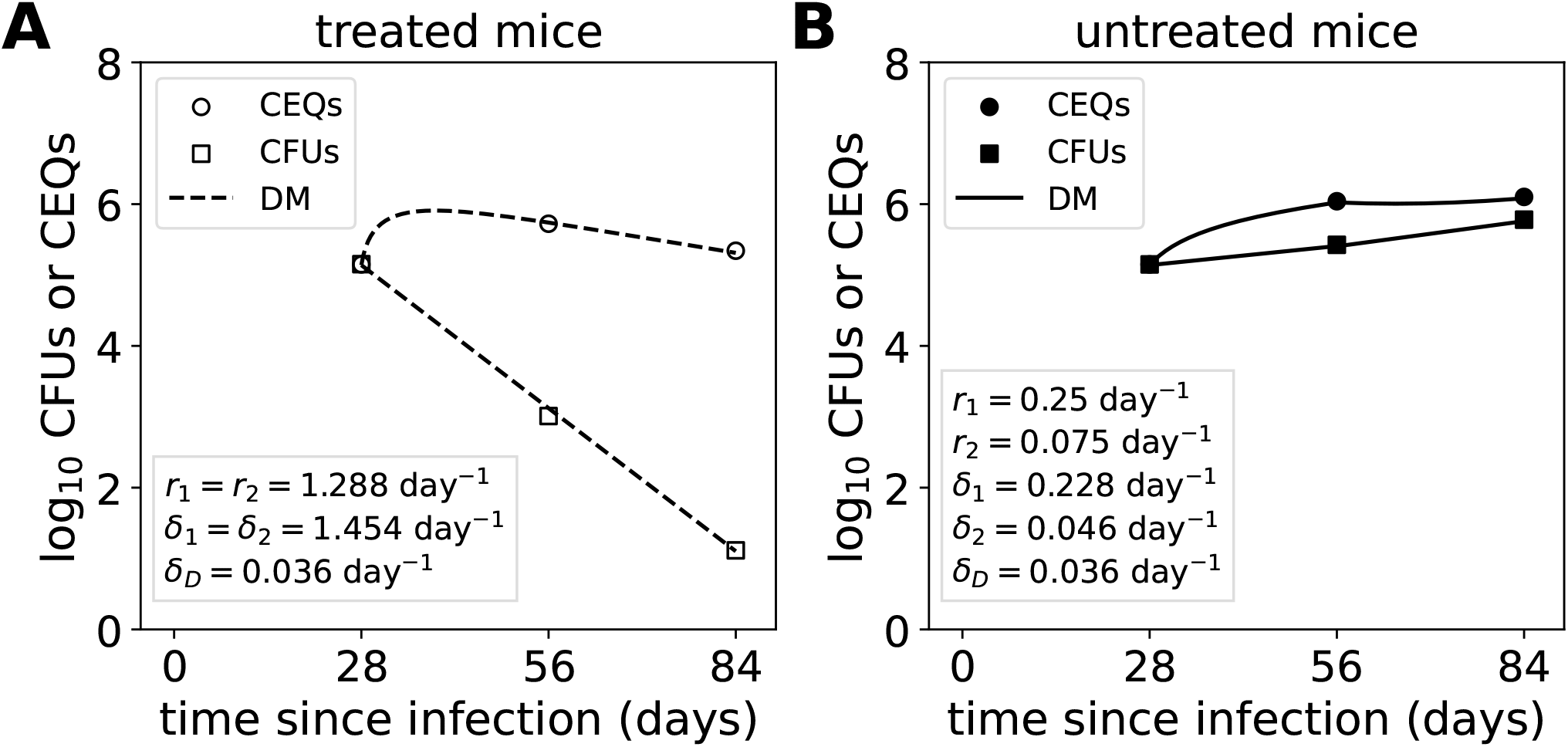
Using DD model to estimate genome decay rate *δ*_*D*_ from CFU and CEQ data in mice. We fitted DD model (**eqns. (8)–(10)**) to CFU and CEQ lung data in INH-treated (**A**) or untreated (**B**) B6 mice infected with a high dose of Mtb intravenously ^6^. In both panels, rates with subscript 1 or 2 represent the rates during the first (28-56 days) and second (56-84 days) time intervals, respectively; note that these intervals are different from those given in **eqn. (13)**. In panel **A**, only one set of rates was used, since best fits with two rates produced replication and death rates that differed by less than 2% between the two time intervals. Note that a large replication rate was required to fit the substantial rise in CEQs between 28 and 56 days post-infection, for treated mice. We initially fit the data with all parameters being free; we found that the best fit produced *δ*_*D*_ = 0.035 day^−1^. In panel **B**, we fixed the CEQ decay rate in the DD model to *δ*_*D*_ = 0.036*/*day and estimated other model parameters are listed on the panel.

**Supplemental Figure S3:**
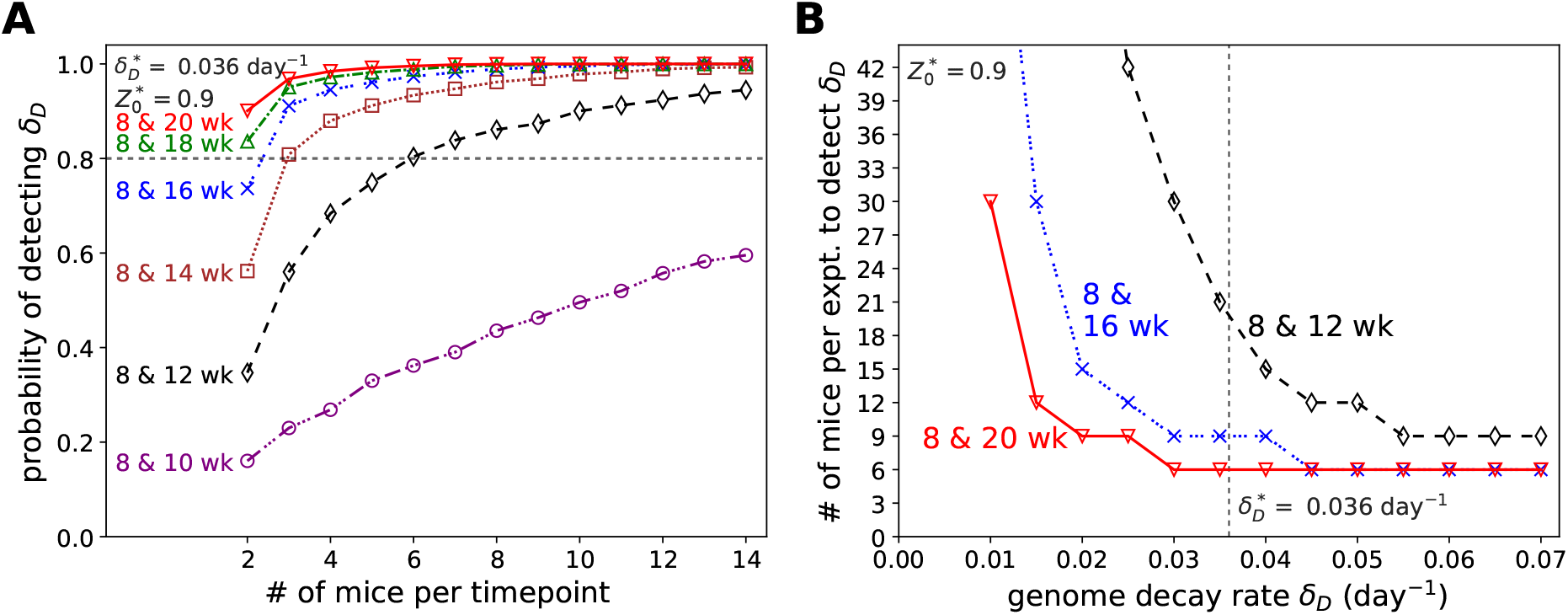
Power analysis to detect Mtb genome decay rate *δ*_*D*_. Similar analyses as in Figure 4 except *Z*(28) = 0.9.

**Supplemental Figure S4:**
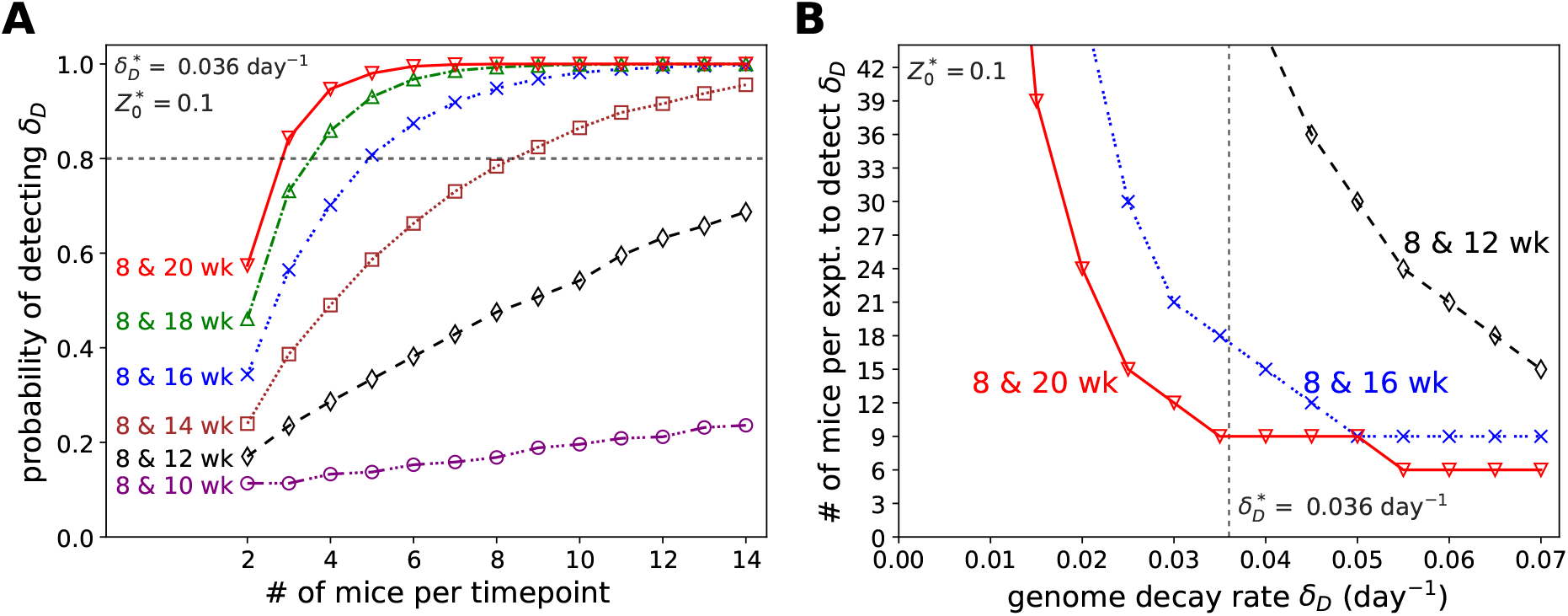
Power analysis to detect Mtb genome decay rate *δ*_*D*_. Similar analyses as in Figure 4 except *Z*(28) = 0.1.

**Supplemental Figure S5:**
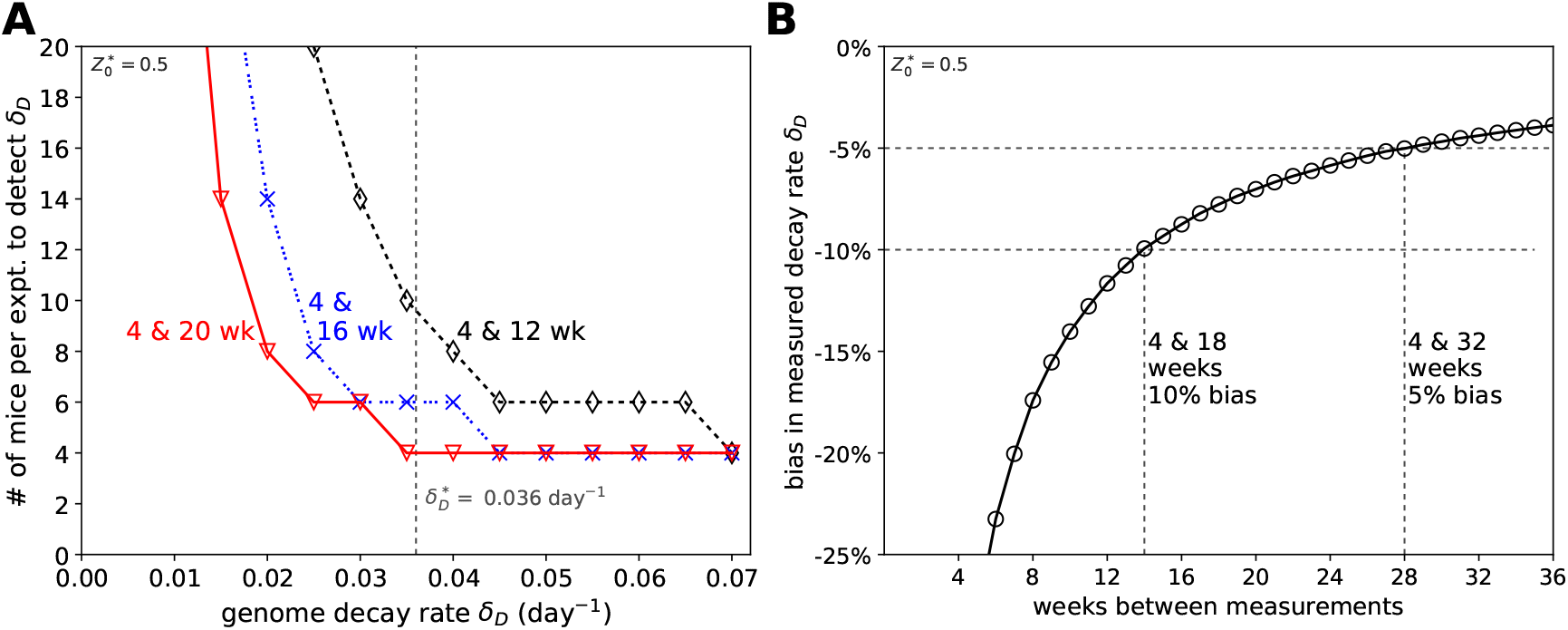
An alternative experiment has higher statistical power but at the cost of introducing bias at estimating Mtb genome decay rate. In this alternative experiment, *B* and *Q* are measured only twice: the first is 28 days post-infection, before antibiotic treatment starts, and the second is at the end of the experiment; thus, this set-up requires 2*n* mice (see **Figure 4**A for comparison). **A**: Number of mice needed for 80% statistical power to detect genome decay rate of *δ*_*D*_ = 0.036*/*day. **B**: Estimated percent bias predicted in measurement of *δ*_*D*_. We estimated the bias by simulating the system with the DD model, with *δ*_*D*_ = 0.036 day^−1^. In simulations we assume that *Z*(28) = 0.5.

**Supplemental Figure S6:**
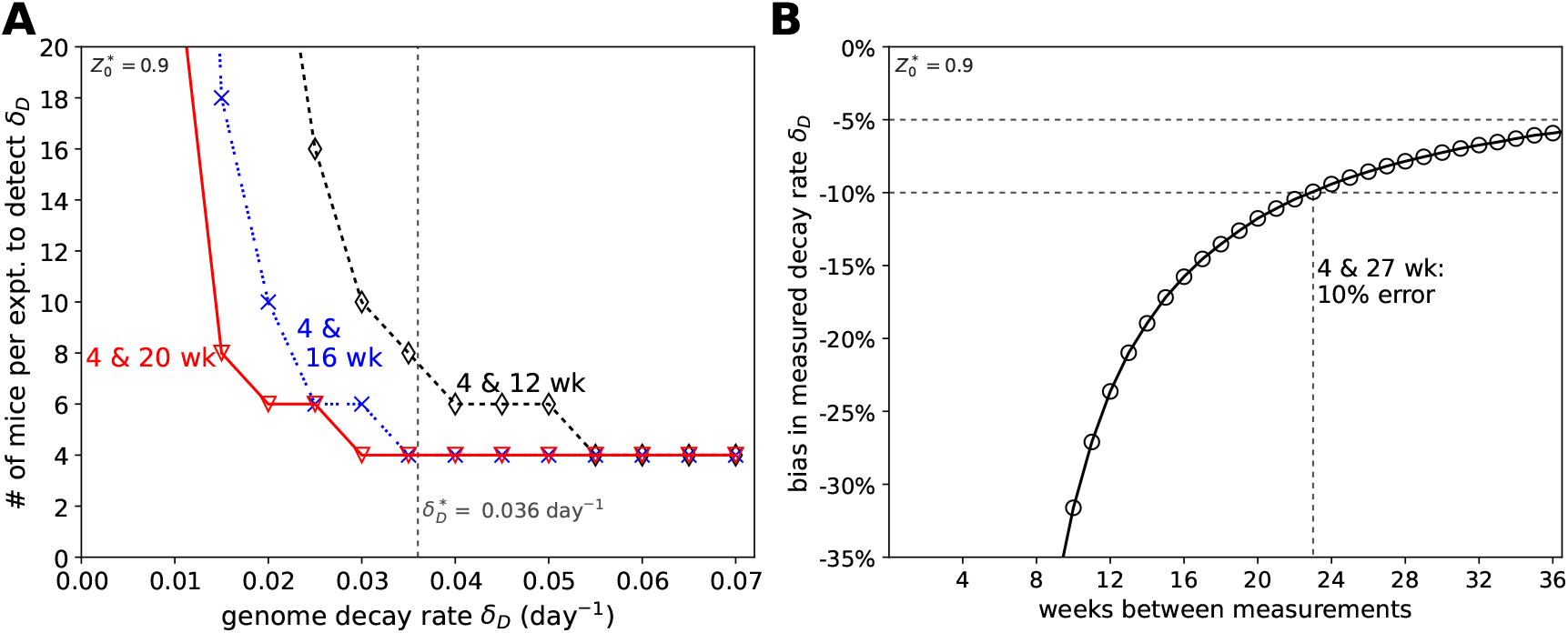
An alternative experiment has higher statistical power but at the cost of introducing bias. Similar as in **Supplemental Figure S5** except we assume *Z*(28) = 0.9.

**Supplemental Figure S7:**
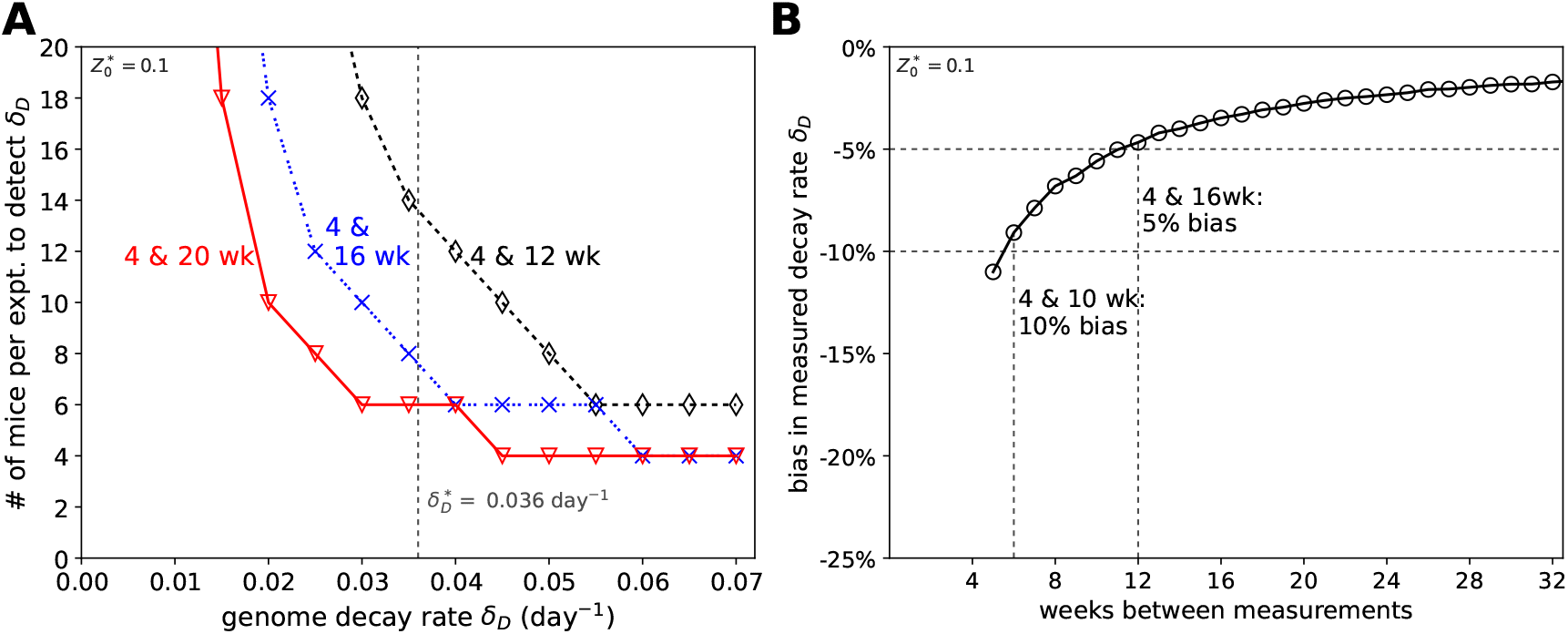
An alternative experiment has higher statistical power but at the cost of introducing bias. Similar as in **Supplemental Figure S5** except we assume *Z*(28) = 0.1.

**Supplemental Figure S8:**
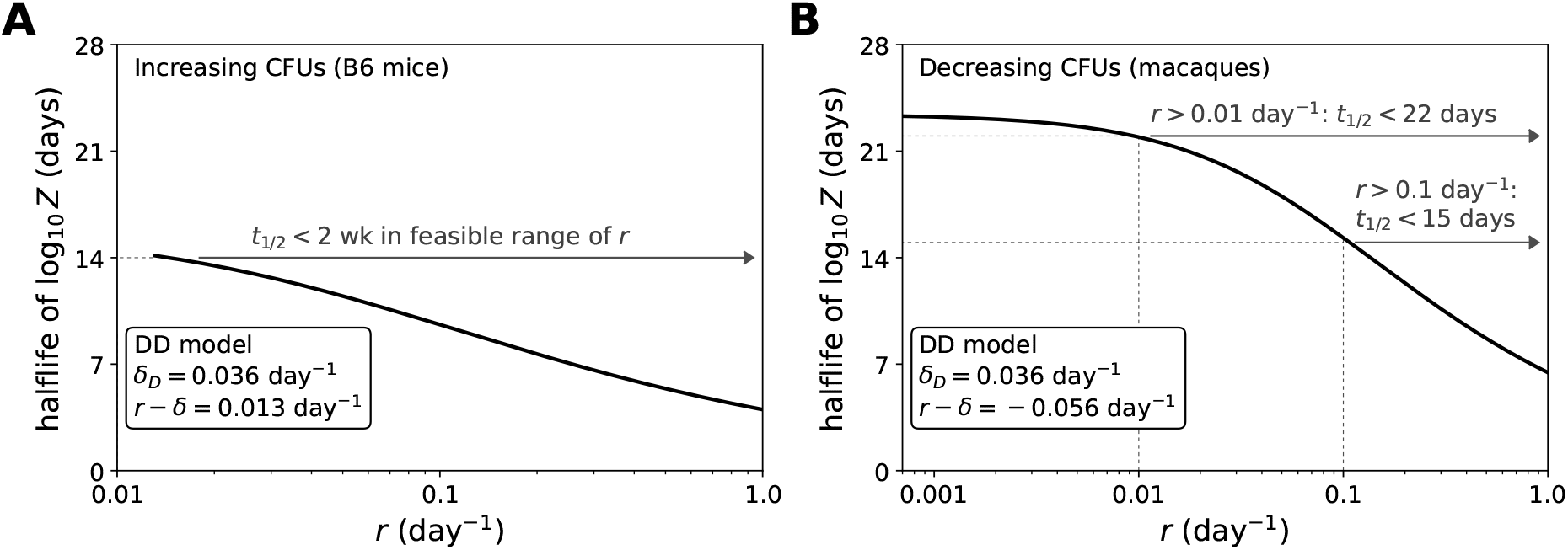
Relatively rapid decay of detectable chromosomes results in a short memory of the CFU/CEQ ratio, when bacterial burdens are slowly increasing. We used the DD model to estimate the timescale over which dynamics of *Z* approach asymptotic behavior. **A**: To find the time for the difference log *Z* − log *Z*_∞_ to decay to half its initial value in mice between 4 and 8 weeks post-infection, we solved the equation log *Z*(*t*) − log 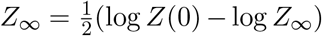 for *t*, where *Z*(*t*) is given by **eqn. (S.13)**, and *Z*_∞_ = 1−*δ/*(*r* +*δ*_*D*_). We assumed that *Z*(0) = 1, and that *r* −*δ* = 0.013 day^−1^, the net growth rate we estimate in mice between 28 and 56 days post-infection (**Table 1**). **B**: To consider the timescale when the population is declining, we found the time for log *Z* − log *Z*_*L*_ to decay to half its initial value, by iteratively solving log *Z*(*t*) − log 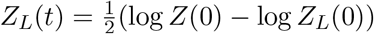 for *t*, where *Z*_*L*_ is given by **eqn. (S.14)**. We again used *Z*(0) = 1, and we set *r* − *δ* = −0.055 day^−1^, the net rate of population decline Mtb in macaques, between 28 and 77 days post infection (**Table 2**).

